# Accurate Conformation Sampling via Protein Structural Diffusion

**DOI:** 10.1101/2024.05.20.594916

**Authors:** Jiahao Fan, Ziyao Li, Eric Alcaide, Guolin Ke, Huaqing Huang, E Weinan

## Abstract

Accurately sampling of protein conformations is pivotal for advances in biology and medicine. Although there have been tremendous progress in protein structure prediction in recent years due to deep learning, models that can predict the different stable conformations of proteins with high accuracy and structural validity are still lacking. Here, we introduce UFConf, a cutting-edge approach designed for robust sampling of diverse protein conformations based solely on amino acid sequences. This method transforms AlphaFold2 into a diffusion model by implementing a conformation-based diffusion process and adapting the architecture to process diffused inputs effectively. To counteract the inherent conformational bias in the Protein Data Bank, we developed a novel hierarchical reweighting protocol based on structural clustering. Our evaluations demonstrate that UFConf out-performs existing methods in terms of successful sampling and structural validity. The comparisons with long time molecular dynamics show that UFConf can overcome the energy barrier existing in molecular dynamics simulations and perform more efficient sampling. Furthermore, We showcase UFConf’s utility in drug discovery through its application in neural protein-ligand docking. In a blind test, it accurately predicted a novel protein-ligand complex, underscoring its potential to impact real-world biological research. Additionally, we present other modes of sampling using UFConf, including partial sampling with fixed motif, langevin dynamics and structural interpolation.

## Introduction

Proteins rely on their three-dimensional structures to participate in biological processes, from catalyzing metabolic reactions to mediating cellular signaling. Recent deep learning advancements, epitomized by AlphaFold2,^1^ have revolutionized protein structure prediction. However, proteins are not static in vivo, they dynamically adopt multiple conformations, enabling interactions with diverse binding partners across distinct states. And *sampling diverse, low-energy conformations of proteins* remains an open challenge. Advanced computational approaches to capture this structural heterogeneity hold profound implications for both biological research and medical applications. As an example, through conformational sampling, researchers can identify transient structural pockets on the surfaces of proteins, ^2^ revealing potential targets for therapeutic interventions that are not discernible in static structural representations. Traditional simulation methods mainly rely on molecular dynamics (MD), ^3^ yet they struggle with the protein’s intrinsic complexity and high free energy barriers, making it difficult to capture essential dynamics on realistic timescales. While enhanced sampling methods have made strides, ^4–6^ the timescale required to capture the essential dynamics and conformational changes is still hard to achieve in general. Recent studies show that with smaller subsampled mutiple sequence alignments (MSAs) or selective MSA clusters as inputs, AlphaFold2 is capable of yielding alternate protein conformations.^7,8^ However, the potential bias introduced by the targeted MSA reduction is yet to be studied. For instance, we find in our experiments that such methods either cannot sample the known structures or do not have high sampling precision. Typical results from MSA clustering, or MSA subsampling with shallow MSA, are that a wide variety of conformations are generated, yet the majority are distant from any known structures. Although the pLDDT metric can be used as a confidence score to select the useful conformations, thermodynamically disfavoured fold-switching (FS) state generated by MSA clustering was found to have a higher pLDDT than the ground state.^8^ So we argue that a good sampler should mainly sample the low-energy conformations.

An alternative approach to conformation sampling is through generative modeling. Unlike MSA subsampling or MSA clustering methods which manipulate the input on trained AlphaFold2 model, generative modeling performs training on resolved protein structures and uses the trained model to sample structures in an unbiased way with only amino acid sequence and noise as input. The rationale behind generative models is that distinct resolved structures with the same sequence exist in the training set, so a structural probability distribution on amino acid sequences is expected to be learned by the model. Unlike molecular dynamics (MD) methods, which inherently perform correlated sampling, generative models facilitate direct sampling after training, thereby circumventing the free energy barriers characteristic of MD techniques. Recent investigations have explored the potential of diffusion- or flow-based models combined with AlphaFold2 to sample a diverse range of protein conformations. ^9,10^ Also recently the original team who developed AlphaFold2 produced AlphaFold3^11^ which incorporates diffusion module in the model. Despite the innovative nature of this approach, our findings indicate that current implementations fall short in terms of sampling accuracy or structural validity. Additionally, it is important to recognize alternative research directions that employ generative models for protein structure generation, primarily focusing on protein design rather than structural sampling.^12,13^

To evaluate the performance of different sampling methods, we hypothesized that chains with rather different structures but with the same sequence in the PDB are good target cases — 47 chains are selected in our evaluation. The rationale behind the testset is twofold: Firstly although protein chains often coexist with other chains as multimers or with other ligands as complex, the structures they adopt in the complex are generally in low-energy states stabilized by the neighboring chains or ligands. Secondly, in a practical viewpoint, when used as targets for rigid docking, the sampled structures of the chains are better in the conformation existing in the complex rather than in their alone to get better results. Thus rather than sampling different conformations of the chains where they adopt in their alone, we argue that a good sampling method could sample conformations of proteins in low-energy states either in their alone or in the complex they could form.

To fully tap the potential of deep generative models in conformation sampling, we propose UFConf, a structural diffusion model based on fine-tuning AlphaFold2. Specifically, we constructed an explicit diffusion process over each protein residue’s position and rotation angle space as in Fig. 1a, and use a modified AlphaFold2 model to predict protein structures given diffused structures at different noise scale as input. As shown in Fig. 1b, we introduce several new modules and modifications to AlphaFold2, both to fully utilize the pre-trained parameters and to accommodate the diffused structural inputs. Briefly speaking, we remove the structure template channel in the AlphaFold2 model, and add a channel to receive the time embedding representing the noise scale. We reuse the channel receiving the initial structure in the structural module in AlphaFold2 model, to receive the diffused structures in UFConf, the details of the model architecture can be found in the Method section. UFConf is trained with continuous noise scheduling, which allows sampling with an arbitrary number of diffusion steps during inference, thus introducing a natural tradeoff between conformation quality and running time. During inference, a noisy structure is sampled randomly, and UFConf is run iteratively to refine the noisy structures to the predicted structures. Additionally, we employ a novel random span-masking strategy during training, which enables partial sampling of conformations with fixed motifs. Most critically, we develop a new hierarchical reweighting protocol of training samples based on structural clustering, which significantly mitigates the bias of conformation distributions in the Protein Data Bank (PDB).^14^

**Figure 1.**
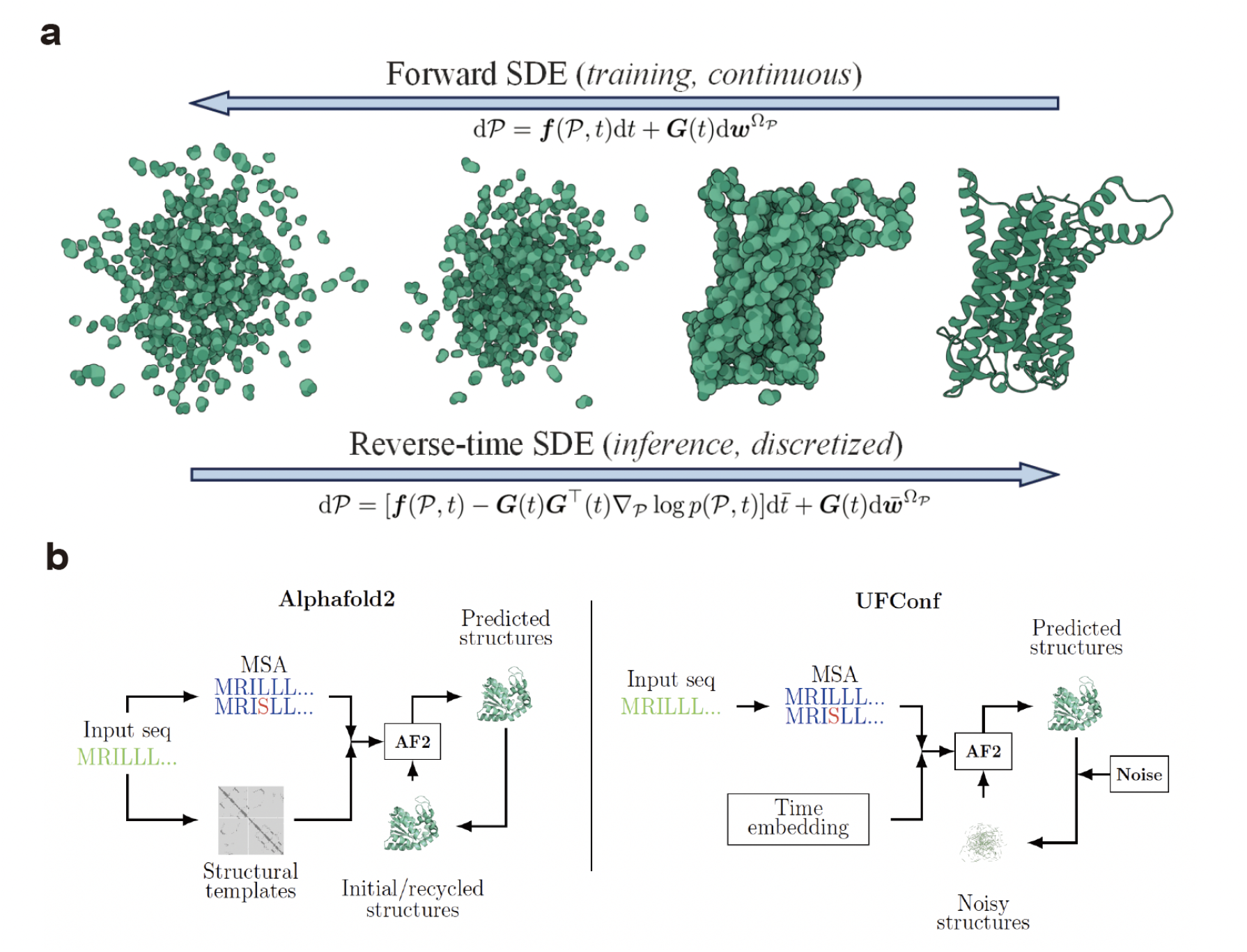
Illustration of structural diffusion and overview of UFConf architecture.(a) Forward and backward structural diffusion process for proteins. During the training phase, UFConf is trained to recover the true structures with different scale of noise added on. During the inference phase, UFConf is run iteratively by reversing the diffusion process from random noise to produce structures for a given sequence. (b) UFConf architecture is modified from AlphaFold2, with structural templates channel removed, and time embedding channel added to represent the input noise scale. The initial/recyvled structures channel is reused in UFConf to receive the noisy structures. UFConf is trained by fine-tuning the trained AlphaFold2 model.

When evaluated on the curated benchmark, UFConf outperforms existing methods in terms of successful sampling and structural validity. Further comparisons with MD simulations show that UFConf can overcome the free energy barrier inherited in the MD methods and perform more efficient sampling. We also show through a case study that the conformations generated by UFConf could improve the performance of the docking method in a blind and cross docking setup, successfully predict the final docking pose (≤ 2.0 Å). We also introduce three novel methods of manipulating UFConf beyond direct sampling, namely *Partial sampling with fixed motif, Langevin dynamics* and *structural interpolation*. The capability of UFConf could enable characterization of multiple conformations of unseen proteins, and aid in the discovery of targeted binding molecules.

## Results

### Training and Inference

UFConf was implemented based on the open-source training code of Uni-Fold,^15^ and the model parameters were initialized from the newest pre-trained weights of AlphaFold-Multimer (v3), released on 2022-12-06 (the training date cutoff is 2018-04-30). All protein chains in the Protein Data Bank released before 2022-04-30 were used in fine-tuning, while no AlphaFold-generated structures (i.e. self-distillation samples) were used. We resorted to Uni-Fold^15^ for generated multiple sequence alignments (MSAs), which followed the MSA curation protocol of AlphaFold. We did not use structural templates. It took 16 NVIDIA A100 GPUs with 80GB memory for approximately 84 hours to fine-tune the model, with a batch size of 64 and a total of 20,000 steps. More details of model training are introduced in Supplementary A.5. In the inference stage, we generated MSAs with the online server of MMSeqs2,^16^ which adopted an MSA search protocol described in ColabFold.^17^ Unless otherwise specified, the reverse processes were discretized uniformly to 30 steps.

### Sampling Alternative Conformations

We evaluated the ability of UFConf in sampling alternative protein conformations on a curated benchmarks, namely RAC-47, in comparison with AlphaFold2,^1^ AlphaFold3,^11^ AF-cluster^8^ (an MSA clustering-based method to generate multiple conformations), MSA-subsampling^7^ (different protein conformations are generated by MSA subsampling) and AlphaFlow^10^ (a recent flow-matching model based on AlphaFold). We also compare the results with DiG^9^ (a diffusion-based model using the Graphormer network^18^) through case study, since the authors only disclose the input feature data files for cases in their paper. Furthermore, UFConf is compared with MD simulation on selected cases, where different experimental structures are chosen as starting conformation for MD simulation.

#### Benchmark performance

We evaluated the performance of different methods by curating a new benchmark, **RAC-47**, abbreviated from *47 recent proteins with alternative conformations*. Specifically, we collected all PDB chains released between 2022-04-30 and 2024-03-29 and selected sequence clusters containing these chains with 100% identity and have 2-10 chains in each cluster. We further filtered the clusters with sequence length between 128 and 768, and the maximum RMSD between chains in the clusters > 2 Å, resulting in 47 clusters of chains. The dataset generation process is unbiased and the details can be found in Supplementary B.1. We sampled 100 conformations with UFConf, AlphaFold2,^1^ AlphaFold3,^11^ AlphaFlow,^10^ AF-cluster^8^ and MSA-subsampling^7^ methods for each of the pairs. Specifically, during inference by UFConf, we subsample the MSA with depth 1024 for each new sample, but not for each inference step, which is the same with the default setup for AlphaFlow. For AF-cluster, we select the first 100 MSA clusters (or all clusters if the cluster number is smaller than 100) generated by its open-source implementation, and ran AlphaFold2 on each. For MSA-subsampling, we run inference by subsampling MSA with depth 64, 256, and 1024. The result for subsampling MSA with depth 1024 is the same with the default setup for AlphaFold2. The same raw MSAs are used for all models. For AlphaFold3 we used the online server provided by the AlphaFold3 team to conduct the inference. We reported the best TM-score of the sampled 100 conformations to the 2 real conformations with largest RMSD in each cluster. Moreover, to evaluate both the diversity and accuracy of the generated models, we used the *matchness score* ^19^ of precision and recall (MAT-P / MAT-R) to assess the overall quality of all generated conformations, which was originally designed to measure qualities of molecular conformations. Specifically,

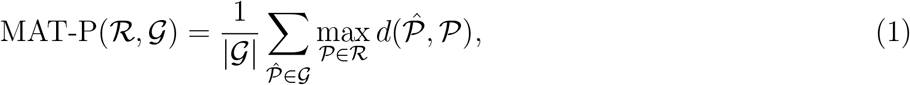

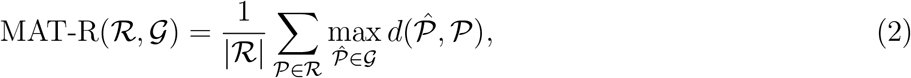

where ℛ and 𝒢 are the sets of real and generated conformations, and we set the metric *d* as the TM-score between two conformations. Intuitively, MAT-P measures the average precision of the generated conformations concerning the closest real conformation, and MAT-R measures how well the real conformations are covered by the generated ones.

Table 1 reports the performance of all models on **RAC-47**. Specifically, we reported the mean and median of the best TM-score among 100 generated conformations corresponding to two real conformations with largest RMSD in each cluster across the 47 cases. Detailed results for individual cases can be seen in Supplementary B.2. From Table 1 we can see that UFConf performs the best in terms of the median TM-score, while AlphaFlow achieves the best mean TM-score. To evaluate different models in terms of how successful the sampling is, we defined a “successful sampling” to be that each of the best TM-score for a given case is larger than 0.90. This is a rational metric considering that the relative TM-score between two experimental conformations of all cases in **RAC-47** ranges from 0.461 to 0.900. From Table 1 we can see that UFConf achieves the most successful sampling with 19 among 47 cases. Additionally, we reported the average MAT-P and MAT-R across the 47 cases. Regarding recovery, UFConf and AlphaFlow performs the best in terms of median and mean MAT-R respectively. In terms of MAT-P, UFConf outperform the MSA-subsampling with MSA depth 64 and AF-cluster, while behind AlphaFold2,AlphaFold3, AlphaFlow and MSA-subsampling with MSA depth 256. The superiority of UFConf can also be seen in Fig. 2a-h, where two typical sampling cases are shown. We could see that compared with AlphaFlow, the sampling of UFConf is more diverse, thus explaining in part why AlphaFlow outperform UFConf in MAT-P. But the diversity of the sampling in UFConf is controlled so that the generated conformations do not deviate from the experimental structures too much, so the diverse sampling exhibited by UFConf might be advantageous in situations where rather different conformations are needed. On the other hand, the diversity of the sampling in MSA-subsampling and AF-cluster are generally uncontrolled, thus leading to mostly unwanted conformations.

**Table 1:** Evaluation results on RAC-47. Mean *Diff*. indicates the mean TM-score between two real conformations across 47 cases. The best TM-score of 100 generated conformations to both of the real ones in each case are reported and aggregated. The depth of MSA for MSA-subsampling model is indicated in the bracket.

| - | Mean *Diff.* | AlphaFold2 | AlphaFold3 | UFConf | AlphaFlow |
| --- | --- | --- | --- | --- | --- |
| Mean-of-Best TM-score (↑) | 0.753 | 0.811/0.783 | 0.831/0.803 | 0.839/0.824 | **0.850/0.826** |
| Median-of-Best TM-score (↑) | 0.757 | 0.866/0.808 | 0.887/0.856 | 0.915/**0.896** | **0.916**/0.876 |
| Num. of successful sampling (↑) | - | 9 | 11 | **19** | 18 |
| mean/median MAT-R (↑) | - | 0.802/0.832 | 0.824/0.872 | 0.838/**0.900** | **0.842**/0.877 |
| mean/median MAT-P (↑) | - | 0.808/0.845 | 0.810/0.866 | 0.768/0.805 | 0.803/0.846 |
| - | MSA-subsampling (64) | MSA-subsampling (256) | AF-cluster |  |  |
| Mean-of-Best TM-score (↑) | 0.818/0.793 | 0.822/0.794 | 0.696/0.688 | | |
| Median-of-Best TM-score (↑) | 0.863/0.809 | 0.856/0.807 | 0.722/0.722 | | |
| Num. of successful sampling (↑) | 11 | 12 | 2 | | |
| mean/median MAT-R (↑) | 0.812/0.845 | 0.813/0.844 | 0.697/0.734 | | |
| mean/median MAT-P (↑) | 0.758/0.777 | 0.802/0.828 | 0.495/0.504 | | |

**Figure 2.**
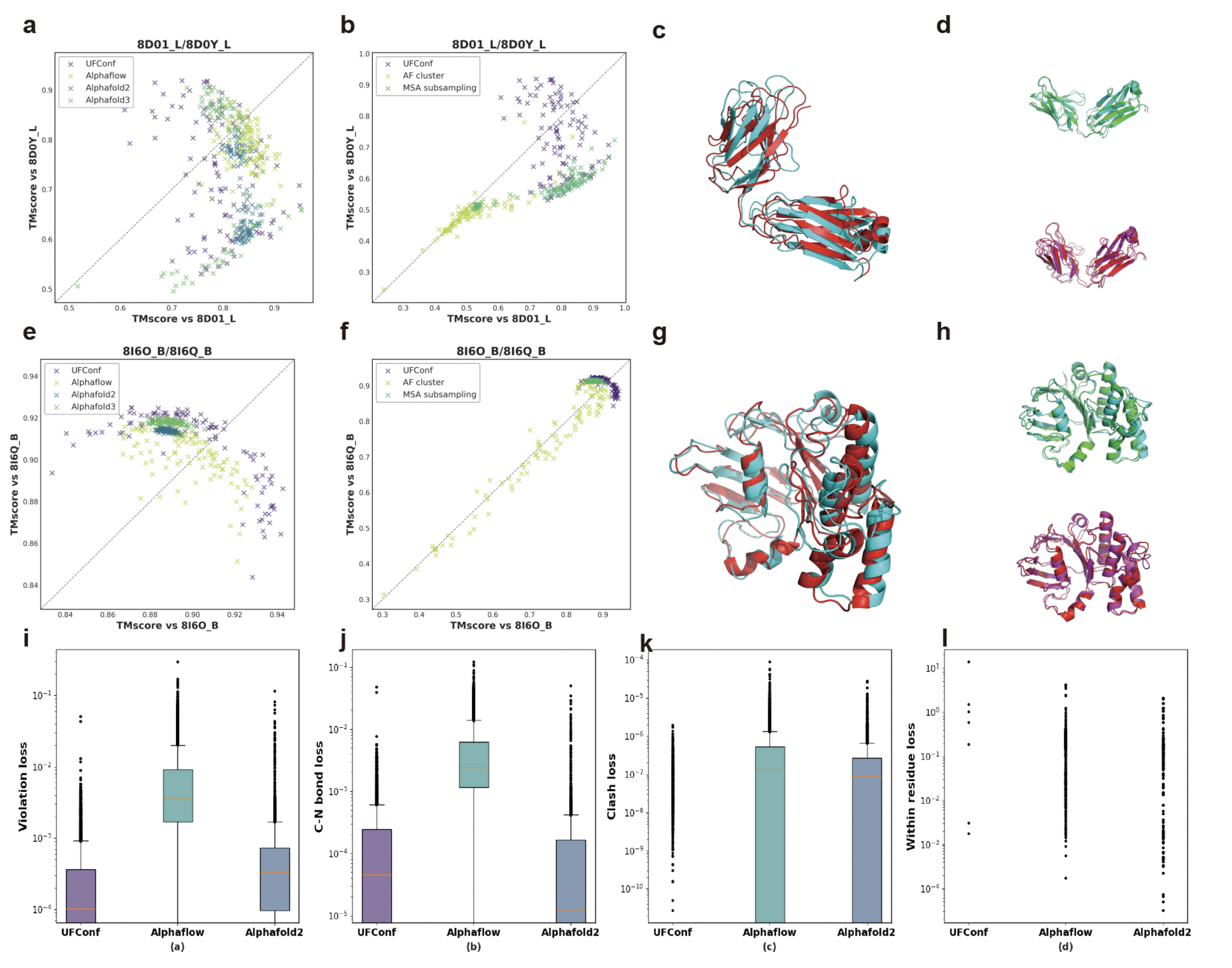
Performance comparison of different models on selected cases and comparison of structural violations between different models. (a)(b) Scatter plot of TM-score between 8D01_L/8DOY_L and 100 conformations generated by six different models. (c)(d) Overlay of sampled structures from UFConf with 8D01_L/8DOY_L. cyan: 8D01_L experimental structure; red: 8DOY_L experimental structure; green: sampled structure closest to 8D01_L; magenta: sampled structure closest to 8DOY_L. (e)(f) Scatter plot of TM-score between 8I6O_B/8I6Q_B and 100 conformations generated by six different models. (g)(h) Overlay of sampled structures from UFConf with 8I6O_B/8I6Q_B. cyan: 8I6O_B experimental structure; red: 8I6Q_B experimental structure; green: sampled structure closest to 8I6O_B; magenta: sampled structure closest to 8I6Q_B.(i) Total violation loss as defined in^20^ for all generated conformations; (j) Carbon-Nitrogen(C-N) bond loss (indicating the violation from the C-N bond length) for all generated conformations; (k) Clash counts between residues (indicating the violations from the atomic radius restriction between residues) for all generated conformations; (l) Clash counts within residues (indicating the violations from the atomic radius restriction within residues) for all generated conformations;

From the above analysis we can see that UFConf outperform MSA-subsampling and AF-cluster methods in terms of sampling recall and displaying reasonable sampling precision, while having comparable performance with AlphaFlow. To further evaluate these two models, it should be noted that structural validity must be taken account. To evaluate whether the generated configurations satisfy the physical constraints induced by the physical law, we calculated the violation loss and different contributions to it according to the established work^1,20^ for the generated conformations across 47 cases in RAC-47 for the best two models so far: UFConf and AlphaFlow. The results from Alphafold was also calculated as comparisons.

As shown in Fig. 2i-l, UFConf exhibited an equal or superior performance with Alphafold on both the total violation loss metric and also different contributions to it, including the C-N bond loss, clash counts between residues, and the clash counts within residues. The clash counts is defined to be the number of atom pairs whose distance is smaller than the atomic van der waals radius with a tolerance. On the other hand, the violation loss and each contributed loss in AlphaFlow are significantly higher than those of UFConf and Alphafold, which indicated that the conformations generated by AlphaFlow have physical invalidity.

#### Comparison with DiG and MD

To compare UFConf with DiG,^9^ we selected cases described in their papers since they only disclose the input files for those cases in their open-sourced code. As shown in Fig. 3a-d, we can see that the sampling of DiG is concentrated on a small region for both cases, while UFConf have a balanced sampling between 1AKE_A and 4AKE_A conformations, and have a more diverse sampling on 2DRI_A/1URP_A case than DiG. Furthermore, it should be noted that conformations generated by DiG only contain the backbone of the proteins without side chains, while all other prediction methods produce the all-atom protein structures.

**Figure 3.**
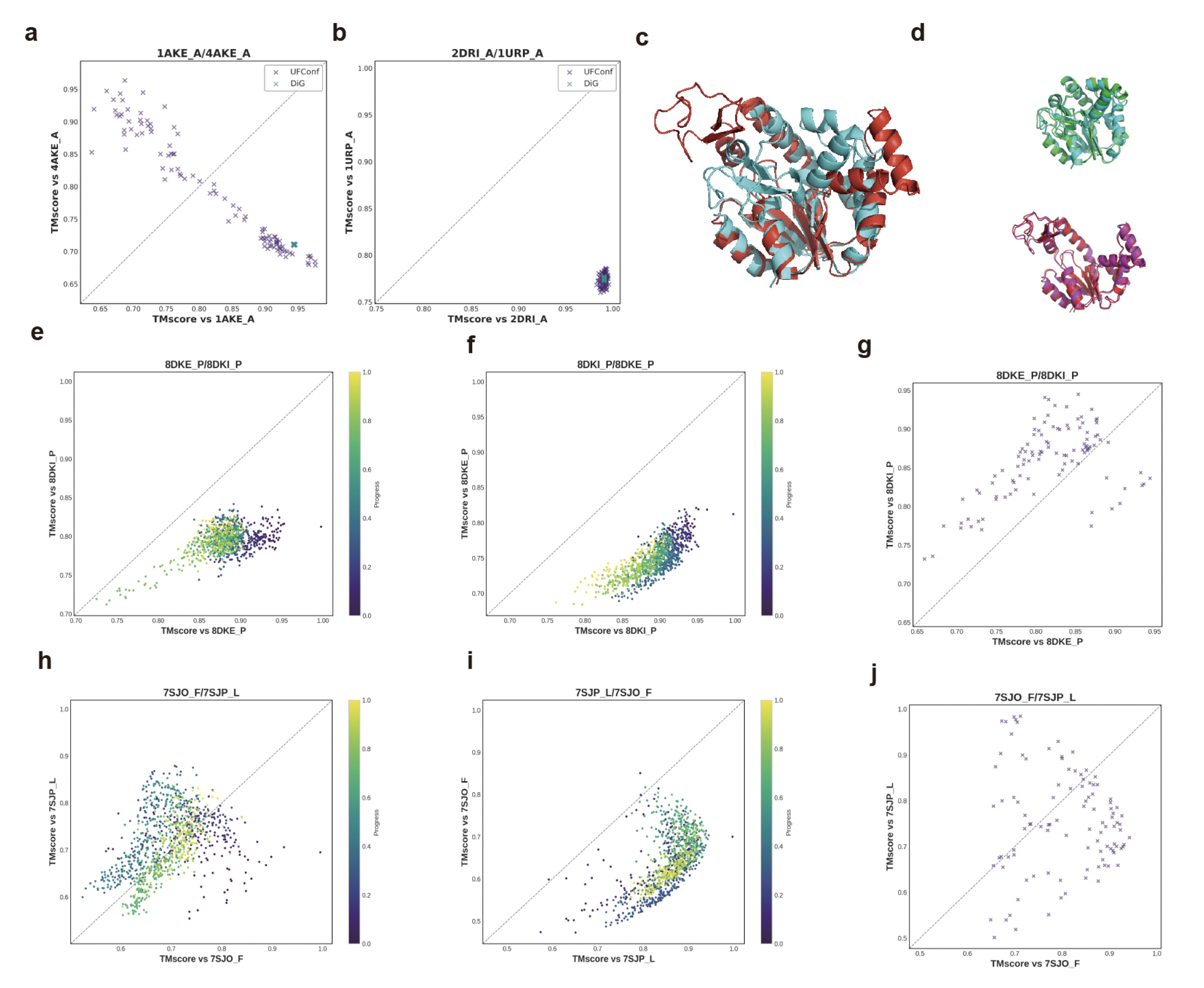
Performance comparisons of UFConf with DiG and with MD simulations. (a) Scatter plot of TM-score between 1AKE_A/4AKE_A and 100 conformations generated by UFConf and DiG. (b) Scatter plot of TM-score between 2DRI_A/1URP_A and 100 conformations generated by UFConf and DiG. (c)(d) Overlay of sampled structures from UFConf with 1AKE_A/4AKE_A. cyan: 1AKE_A experimental structure; red: 4AKE_A experimental structure; green: sampled structure closest to 1AKE_A; magenta: sampled structure closest to 4AKE_A. (e-f) Scatter plot of TM-score between 8DKE_P/8DKI_P and 100 conformations generated by MD simulations, where the MD simulations are start from 8DKE_P in (e), and start from 8DKI_P in (f), the colorbar indicates the simulation progress. (g) Scatter plot of TM-score between 8DKE_P/8DKI_P and 100 conformations generated by UFConf.(h-i) Scatter plot of TM-score between 7SJO_F/7SJP_L and 100 conformations generated by MD simulations, where the MD simulations are start from 7SJO_F in (h), and start from 7SJP_L in (i). Scatter plot of TM-score between 7SJO_F/7SJP_L and 100 conformations generated by UFConf.

Besides comparing UFConf with different machine learning based methods, we present a comparative analysis of our model UFConf against results from molecular dynamics (MD) simulations. The MD simulations were conducted over a duration of 100 ns. Due to the substantial computational demands associated with prolonged MD simulations, we limit our comparison to selected cases. As illustrated in Fig. 3e-j, the MD simulations fail to explore alternative conformations from a starting conformation in the 8DKE_P/8DKI_P pairs, whereas UFConf successfully samples both conformations within 100 samples. Notably, while MD simulations can transition from the 7SJO_F to the 7SJP_L conformation, they do not reciprocate this transition, suggesting a potential stability preference for the 7SJP_L conformation. In contrast, UFConf demonstrates the capability to sample both conformations effectively within 100 iterations.

#### Ablation study

To assess the impact of our proposed hierarchical reweighting protocol based on structural clustering in the training dataset, and also the discretization steps during inference, we trained an additional version of UFConf using AlphaFold’s sequence clustering-based sampling schema, where sample weights were determined by the inverse sequence cluster size and conformations with identical sequences were uniformly sampled. All experiments were conducted on RAC-47, with the mean MAT-P and MAT-R results summarized in Table 2. We find that the model can produce fairly good predictions with even 1 step from the prior. The precision of the model was impacted with 10 steps, as errors were not negligible under such coarse discretizations. Adding inference steps can improve recall performance, though gains marginalize after 30 steps (whose results are reported as default). Removing the hierarchical reweighting protocol leads to degraded precision and recall. The results corroborate the importance of the reweighting protocol.

**Table 2:** Impacts of the number of inference steps and the hierarchical reweighting protocol on RAC-47.

| #samples | Reweighting | #steps | mean MAT-P (↑) | mean MAT-R (↑) |
| --- | --- | --- | --- | --- |
| 100 | Yes | 1 | 0.767(+0.07) | 0.815(-0.17) |
|  | Yes | 10 | 0.750(-0.10) | 0.820(-0.12) |
|  | Yes | 30 | 0.760(+0.00) | 0.832(+0.00) |
|  | Yes | 50 | 0.760(+0.00) | 0.831(-0.01) |
|  | No | 30 | 0.722(-0.38) | 0.801(-0.31) |
| 5 | Yes | 30 | 0.755(+0.00) | 0.772(+0.00) |
|  | No | 30 | 0.727(-0.28) | 0.733(-0.39) |

### Case Study

Recent research^21^ has evidenced the dynamic nature of SLC15A4 and the crucial role its multiple conformations play in the TLR-IRF5 immune signaling, whose misregulation thought to be crucial in *systemic lupus erythematosus* (SLE) and related diseases. Specifically, the TASL binds to SLC15A4 in its cytoplasmic inward-open confromation (PDB ID: 8JZU), participating in the nucleic acid-sensing TLR7/8 pathway activating IRF5. While the newly characterized inhibitory molecule *feeblin* acting as an allosteric inducer, stablizes SLC15A4 in a TASL binding-incompetent lysosomal outward-open conformation (PDB ID: 8JZX), leading to degradation of TASL. To showcase the usage of conformation sampling for accurate modeling of multi-state proteins, we apply UFConf in a blind setup to SLC15A4 and show its importance for accurately characterizing the newly discovered inhibitor-binding mode. It is important to note that the two experimental conformations are out of the training set of UFConf.

Before demonstrating our method, it should be noted that in a easier redocking setup where the experimental structure (8JZX) is used as the target for docking, existing well-known docking methods like quickvina-W, smina and diffdock^22–24^ can not find the correct (ligand RMSD ≤ 2.0 Å) docking pose, the details are given in Supplementary B.3. We now turn to illustrate our method in a more challenging setup where the protein structure information is completely unknown.

The protocol consisted of sampling 100 conformations with UFConf. From Fig. 4e we can see that both cytosol-open (8JZU) and lysosome-open conformations (8JZX) could be sampled (TMscore > 0.9). Pockets for all conformations were identified with FPocket 4.0^25^ and default parameters and correlations to the TASL binding pocket were investigated. Interestingly, the volume of a central pocket accessible from the lysosome side was found to correlate with protein conformation and inversely with TASL in volume, so it was selected for docking of *feeblin*. A ligand conformation and predicted confidence (pRMSD) were produced for each conformation using the UniMol Docking and UniDock tools. ^26,27^ Prediction confidence was smoothed by KNN (K=3) over predicted ligand conformers. From Fig. 4f we can see that the predicted RMSD is correlated with the true RMSD of the ligand (*Spearman ρ*(pRMSD, RMSD) = 0.74, *p* < 10^−17^) and with the protein conformations (*Spearman ρ* (pRMSD,TM-8JZU - TM-8JZX) = 0.50, *p* < 10^−6^). The prediction confidence and the actual performance is generally better when the predicted conformations are closer to lysosome-open conformation (8JZX). The best prediction by smoothed confidence was taken, which obtains ≤ 2.0 Å ligand RMSD to the experimental structure (PDB ID: 8JZX).

**Figure 4.**
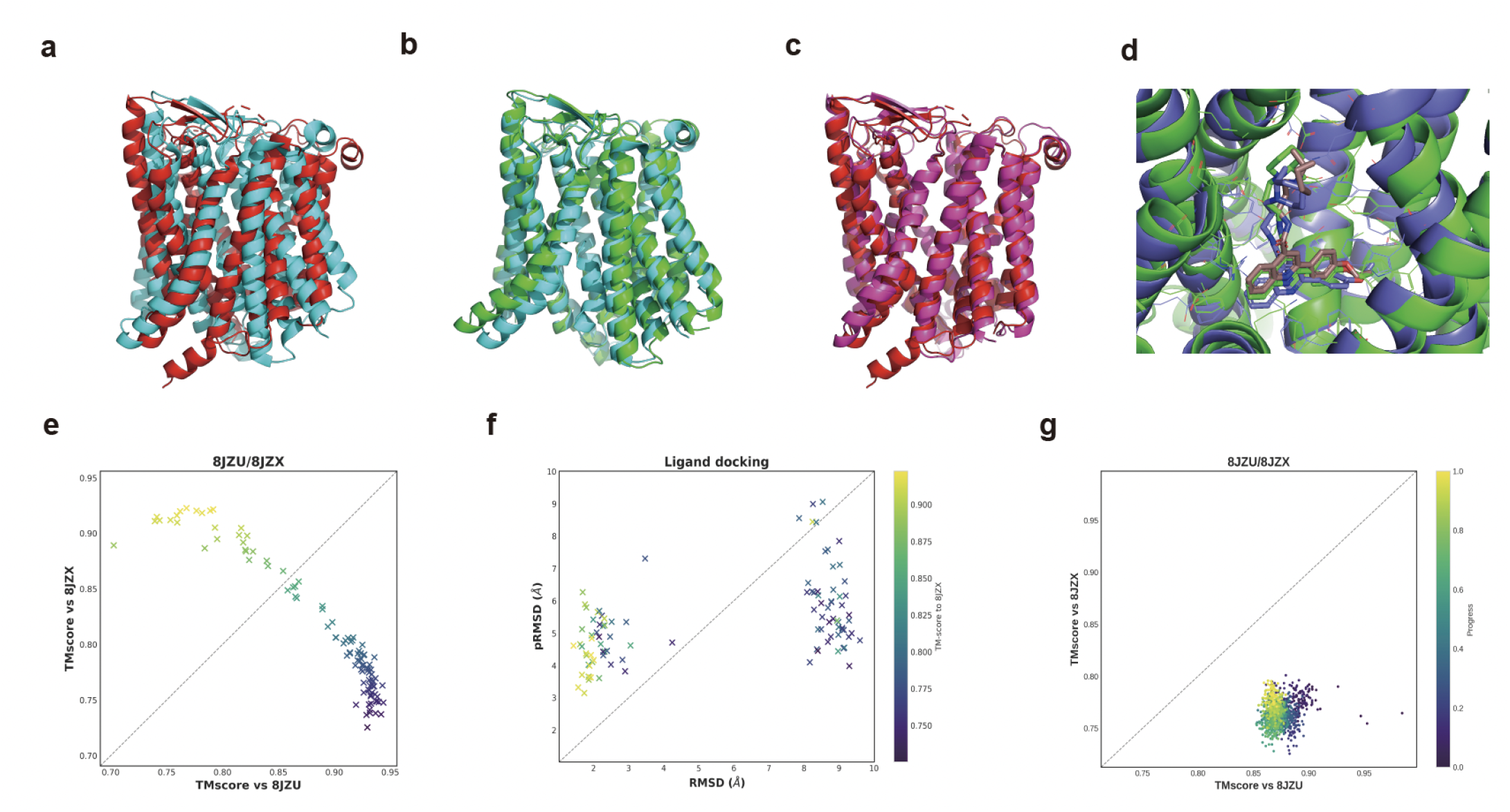
Visualization and scatter plots of SLC15A4 conformations and binding poses. (a-c) Overlay of sampled structures from UFConf with 8JZU/8JZX. cyan: 8JZU experimental structure; red: 8JZX experimental structure; green: sampled structure closest to 8JZU; magenta: sampled structure closest to 8JZX. (d) Visualization of binding site. green: 8jzx experimental structure; brown: ligand bound to experimental structure; blue: predicted ligand and protein structure. (e) Scatter plot of TM-score between the 100 generated conformations and 8JZX/8JZU.(f) Scatter plot of ligand predicted RMSD v.s. RMSD using each sampled conformation, the colorbar indicates the TM-score to 8JZX where the same color corresponds to the same conformation to (e). (g) Scatter plot of TM-score between 8JZU/8JZX and 100 conformations generated by 100ns MD simulations, where the MD simulations are start from 8JZU. The colorbar indicates the simulation progress.

Notably, as shown in Fig. 4g, MD simulation from 8JZU can not sample 8JZX conformation during 100ns run, indicating an energy barrier between the two conformations. This example demonstrates the utility of sampling multiple protein conformations using UFConf to improve the docking performance, thus illuminating the usage of the model in real drug discovery scenarios.

### Partial Sampling, Langevin Dynamics and Structural Interpolation

UFConf can be used to perform partial sampling of proteins with fixed motif, which is illustrated in one example in RAC-47 benchmark. We selected 7UTI_Z/7UTL_d case since the sampling performance from UFConf is bad, as shown in Fig. 5b. Further investigation indicated that it is mainly due to the flexible loop between the two domains in the protein which leads to bad performance. As shown in

**Figure 5.**
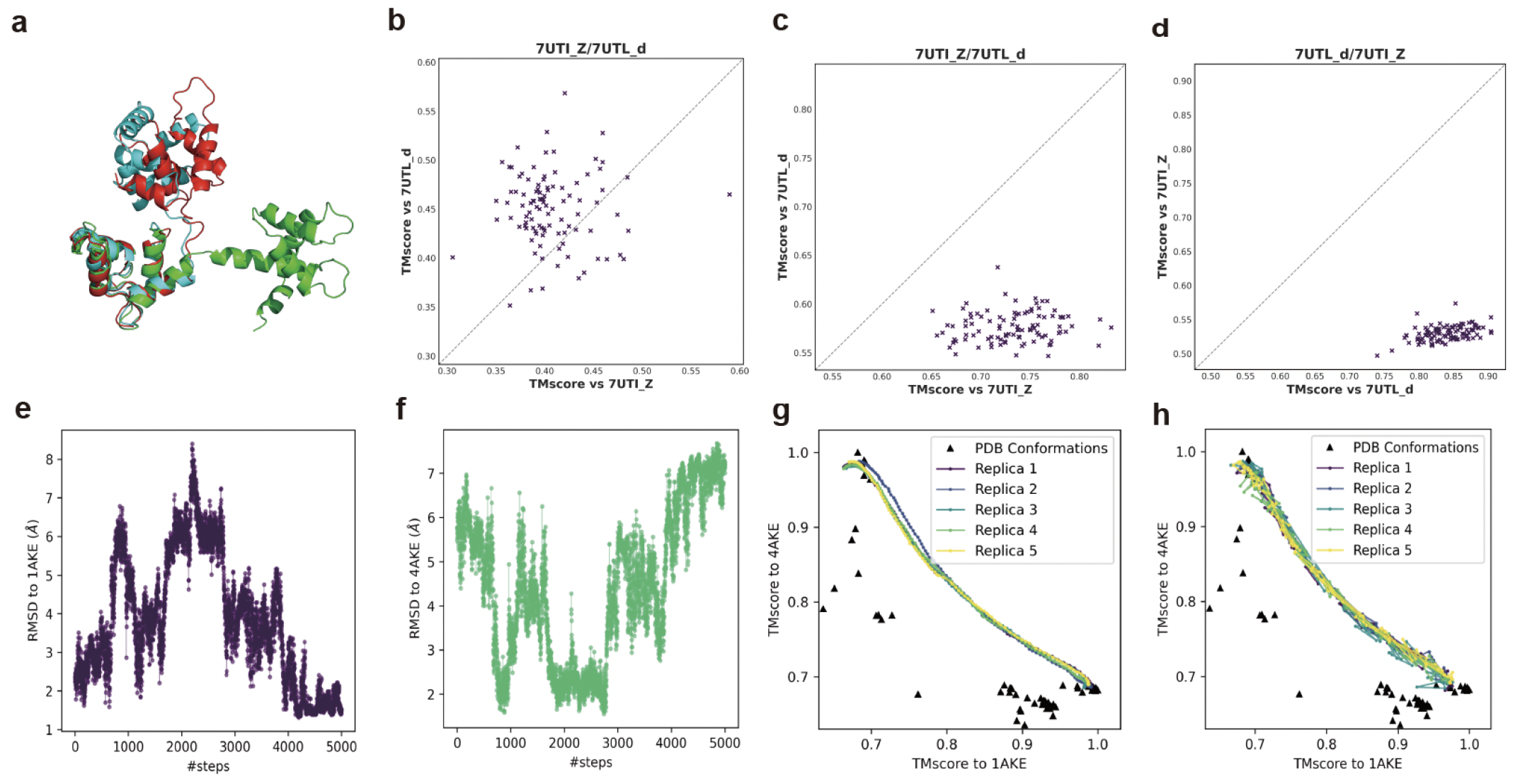
Illustration of partial sampling, langevin dynamics and structural interpolation through case study. (a) Overlay of sampled structures from UFConf with 7UTI_Z/7UTL_d. cyan: 7UTI_Z experimental structure; red: 7UTL_d experimental structure; green: sampled structure from UFConf. (b) Scatter plot of TM-score between 7UTI_Z/7UTL_d and 100 conformations generated by UFConf. (c) Scatter plot of TM-score between 7UTI_Z/7UTL_d and 100 conformations generated by UFConf with left domain and loop region of 7UTI_Z fixed. (d) Scatter plot of TM-score between 7UTI_Z/7UTL_d and 100 conformations generated by UFConf with left domain and loop region of 7UTL_d fixed. (e-f) RMSDs of conformations in Langevin dynamics 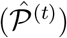 to the two real conformations. (g-h) TMscores of interpolated conformations to the two real conformations, with 1 and 10 inference steps respectively. Other PDB structures in the kinase family are also shown.

Fig. 5a, the middle loop has large diversity, the loop direction of a typical sampled structure (green) from UFConf deviates a lot from the two experimental structure (cray and red), thus leading to large structural disimilarity. We hypothesized that fixing the loop direction aligned with the experimental structures will lead to increase in the sampling accuracy. The results showed that, when using the left domain and the loop region from the 7UTI_Z (cray) experimental structure as fixed motif, the sampled structures will be close to 7UTI_Z with reasonable structural diversity as shown in Fig. 5c. On the other hand, when using the left domain and the loop region from the 7UTL_d (red) experimental structure as fixed motif, the sampled structures will be close to 7UTL_d with reasonable structural diversity as shown in Fig. 5d.

Sometimes we not only want to sample low-energy structures, but also want to see the continuous transition between two stable structures. This could be done using UFConf in the framework of Langevin dynamics and structural interpolation. We illustrate the effects of Langevin dynamics and structural interpolation on adenylate kinase, a classic case of alternative conformations in apo and holo states. This nucleoside monophosphate kinase exhibits an open apo conformation (PDB ID: 4AKE) while adopting a closed conformation (PDB ID: 1AKE) when bound to an inhibitor.^28^ We conduct 5000 steps of Langevin dynamics on the diffused conformations of *t* = 0.6, by iteratively integrating the forward and reverse processes with time step *τ* = 0.01. The starting point was generated with the forward SDE on a predicted structure with *t* = 0.6. For interpolation, we diffused both 1AKE and 4AKE to *t* = 0.3, interpolated the structures with 100 intermediate steps, and predicted results with the reverse-time SDE from *t*, discretized with step size 1 or 10. Fig. 5 visualizes the generated trajectories. From Fig. 5e and Fig. 5f, we observe switches of conformations between 1AKE and 4AKE and several stationary states. More details and discussions are in Supplementary B.5. Fig. 5g and Fig. 5h show that the interpolation results in a putative, continuous transition pathway between the two conformations. Using a discretized inference process of 10 steps (h) leads to more vibration of the structure compared with the direct prediction (g), while the overall trend remains the same.

## Discussions

After the revolutionary breakthrough in protein structure prediction by AlphaFold2, determining different conformations of the proteins is another harder problem with great significance. Success at the problem could deepen our understanding of the dynamic nature of proteins and provide great help in structure based drug discovery. Our results demonstrate that by utilizing the capability of AlphaFold2 and finetuning it to a diffusion model, and through a dataset reweighting protocal based on structural clustering, UFConf could achieve the best performance on curated benchmark RAC-47 among all compared models, successfully sample the most known protein chains with high structural validity.

We also compare UFConf with MD simulation results on selected cases, the results show that UFConf can overcome the energy barrier existing in MD simulations, thus sampling the stable conformations in a more efficient way. There are two possible reasons for the superior sampling efficiency of UFConf over MD simulations: Firstly it might be that it is mainly due to the direct sampling capability of the generative models which overcome the energy barrier existing between different low-energy states. Secondly it might be that it is due to the learned probability distribution itself by UFConf such that there is no free energy barrier or only low barrier existing between different states. Exploration of the possible reason is interesting in its own right and leaves as future work.

We further validated the utility of UFConf through a case study in blind and cross docking setups, demonstrating its ability to sample both cytosol-open and lysosome-open conformations from sequence input alone, thereby enhancing docking outcomes. We could see the whole pipeline of first using UFConf to sample conformations of a protein and then using docking methods to perform rigid docking as a way of doing ensemble docking. Since the prediction confidence of the docking method used could be an indicator to select the best binding pose, the whole pipeline can also be viewed as one way of flexible docking where the targeted protein structure could be changed.

It should be noted that we adopt a different viewpoint for what a good sampling model should be than previous works, where they claim that the sampled structures should be drawn from the boltzmann distribution or assigned a boltzmann weight,^8,9^ characteristic of the canonical probability distribution. Firstly both AlphaFold2 and different generative models (UFConf, DiG, AlphaFlow) are mainly trained on resolved experimental structures, and these structures exist alone or with their binding partner. So the expectation of the trained models is to sample conformations of proteins in their alone or in the complex, unless the training dataset is changed from resolved experimental structures to the conformations from MD simulations. It is reasonable for an isolated protein to conform the canonical distribution, but in combination with the binding partners it is difficult to tell the distribution change of the proteins. So we hypothesize that a good sampling model should sample the conformations of proteins in low-energy states either in their alone or in the complex they could form, without the need to be drawn from a boltzmann distribution or assigned with the boltzmann weight unless there is specific needs to do so. While on the other hands, an indicator should indeed be assigned to the sampled structures as the stability or validity of the conformation, this might be done through calculation of the free energy of different conformations and serves as the future direction.

Another research direction is to extend the framework to protein multimers, whereas the challenge is as well to provide a balanced sampling protocol. Furthermore, we find in our experiment that by performing MSA clustering and using clustered MSA as UFConf input, the model could predict even more diverse structures, but we did not explore more on the combination of generative models and MSA manipulation methods which could also be interesting. We hope that beyond AlphaFold2, methods which could sample different conformations of proteins in low-energy states could provide us more understanding of the protein dynamics and functions.

## Method

### Preliminaries

#### AlphaFold2

AlphaFold2^1,29^ is a deep learning system that accurately predicts protein structures from amino acid sequences, multiple sequence alignments and structural templates. In summary, it introduces *Evoformer*, an attention-based network to encode input sequences and templates, and a *structure module*, an SE(3)-equivariant network to decode the predicted structure step-wise from the origin. A recycling mechanism is introduced to iteratively refine the predicted structure with the same network parameters. Subsequent works^15,20^ implemented open-source training code of AlphaFold2 to enable further development of the model. We implemented UFConf based on.^15^

#### Notations and structure parameterization

We build structural diffusion processes based on the parameterization of protein backbones introduced in AlphaFold2. Formally, a residue 𝒜_*i*_ in a protein structure 𝒫 = (𝒜_1_, *· · ·*, 𝒜_*n*_) is represented by 𝒜_*i*_ = (*T*_*i*_, 𝒳_*i*_), where *T*_*i*_ = (***R***_*i*_, ***c***_*i*_) ∈ SE(3) is the local frame of the residue, defined by the coordinates of the backbone atoms (N,C_*α*_,C), and 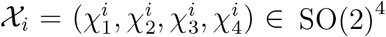 are (up to) four torsion angles which fully determine the flexibility of the sidechain. We use *n* for the length of the protein, *i, j* for residue indices, and 𝒮 = (*s*_1_, *· · ·, s*_*n*_) for the sequence of the protein.

#### Diffusion models

Diffusion models^30,31^ have now become the mainstream method in computer vision. As probabilistic generative models, they characterize data generation as reverse-time diffusion processes. Further research^32^ illuminated deep connections between diffusion models and score-based generative models. In a diffusion model, a stochastic diffusion process is created for ***x*** ∈ Ω_***x***_ as

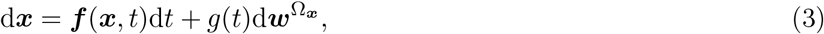

where ***f*** (*·*), *g*(*·*) are pre-defined drift and diffusion factors, and d***w*** is the differential of a standard Wiener process. The reverse-time diffusion process takes the form of

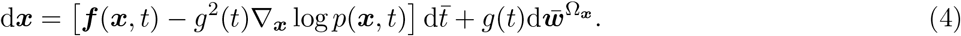

The training of the network aims to learn a parameterized score function ***s***_*θ*_(***x***, *t*) approximating ∇_***x***_ log *p*(***x***, *t*), which is only feasible (conditioned on data samples ***x***_0_ ∈ 𝒟 ⊂ Ω_***x***_) during model training. Recent work has extended diffusion models to protein domain for applications in protein design^12,33,34^ and conformation sampling.^9,35^ When modeling the rotations and torsion angles of proteins, diffusion processes on Riemann manifolds^36^ should be introduced.

#### Diffusion models on Riemann manifolds

Riemannian diffusion models are extensions of Euclidean diffusion models which models the diffusion processes on Riemannian manifolds. In a Riemannian diffusion model, Wiener processes in Eq. (3) and (4) are replaced with Brownian motions on the Riemann manifold ℳ, and score functions ∇_***x***_ log *p*_*t*_(***x***) should be calculated as Riemannian gradients which lies in the tangent space 𝒯_***x***_ℳ. We refer our readers to^36^ and^13^ for theories and more discussions. Our work adopts a similar definition of protein diffusion processes to,^12^ where the main differences are that torsion angles are incorporated and the forward process is not discretized during training.

### Structural Diffusion

Fig. 1a illustrate the concept of structural diffusion. Overall, we create a diffusion process over the protein backbone space Ω_𝒫_ = {SE(3)}^*n*^, and the components on position and rotation angles of each protein residue are defined independently. For simplicity, we hereby describe the diffusion process for a specific residue 𝒜 ∈ 𝒫. For each backbone pose *T* = (***R, c***), we formulate an Euclidean diffusion process for the position and an SO(3) diffusion process for the rotation:

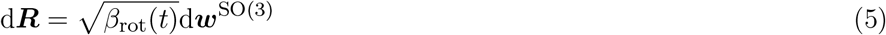

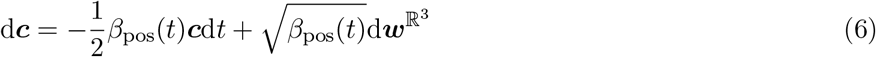

where d***w***^*^s are differentials of Brownian motions in the corresponding spaces, and *β*_*_s are pre-defined functions over *t*. Correspondingly, the marginal density of (***R, c***) at *t*, conditioned on (***R***^(0)^, ***c***^(0)^), can be derived following^32^ as

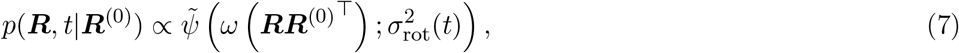

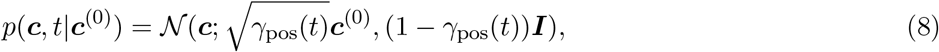

where 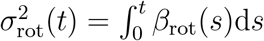 and 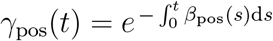 are integrated from *β*(*t*). Here *ω*: SO(3) → [0, *π*) maps an SO(3) element to a rotation angle, and *ψ*(*·*) is the density function of rotation angles of isotropic Gaussian distribution on the SO(3) manifold (IGSO(3), Nikolayev et al. ^37^), defined as

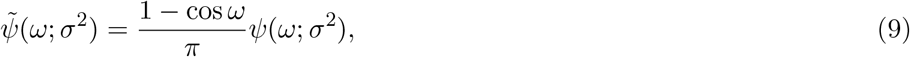

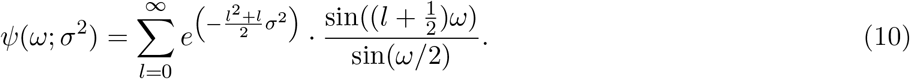

The difference between 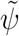 and *ψ* is up to the volume density on the SO(3) manifold with regard to its Haar measure. The forms of the score functions can then be derived as

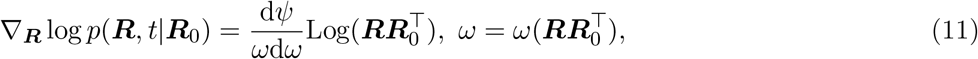

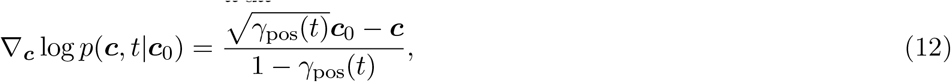

where *ψ* and d*ψ/*d*ω* can be approximated numerically by truncating the summation to *l* ≤ *L*.

The torsion angles are trained and generated in the same way with AlphaFold2^1^ though a shallow ResNet, thus do not undergo diffusion process. We tried to implement diffusion process on torsion angles, but the results were worse than the original model, this might be a research direction for future study.

In choosing the parameters of the diffusion process, we scaled the process in a continuous unit time interval *t* ∈ [0, 1]. We set a distorted *cosine schedule* ^38^ for the position diffusion and polynomial schedules for rotation and torsion diffusions. Specifically,

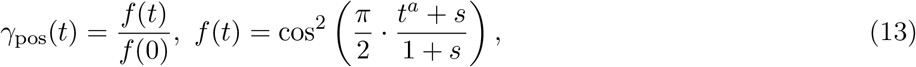

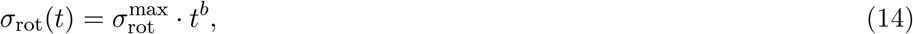

where *a, b* are chosen to empirically smooth the variance along the diffusion process. In positional diffusion, the center-of-mass of the protein is removed and the coordinates are normalized to 1*/*20.

Notably, instead of directly producing the estimated score functions, our model reparameterizes the score function by predicting 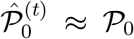.The estimated score function is then calculated by replacing elements in 𝒫_0_ in the above equations with those in 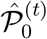. This approach allows the utilization of intermediate model outputs as sampled protein conformations, as well as direct supervision of model outputs to enforce correct geometry. Detailed algorithms for model training and inference are provided in Supplementary A.4.

To further allow partial sampling of protein conformations where a portion of the structure (motif) is fixed, we sample continuous motifs and fix them during training. Specifically, we propose a Markovian random span-masking strategy that generates motif masks with expected continuous segment size *K* and mask probability *ρ*. To generate a sequence of binary masks (*M*_1_, *M*_2_, · · ·), we create

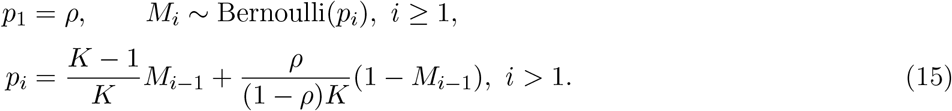

Properties of this span-masking are proved in Supplementary A.1.

### Architecture Modification

The motivation of all following modifications to the architecture of AlphaFold2 is to appropriately transform the protein folding model, *m*_*θ*_(𝒮), to a diffusion model, *m*_*θ*_(*t, P*^(*t*)^, 𝒮). To achieve this, we introduced several new modules to Evoformer and altered the initial structure inputs to the *structure module* of AlphaFold2. We shut the template channel of AlphaFold2, as we wanted the input diffused structures to be the only source of structure information, thus dominating the outcomes of the model. A total of 1.6M trainable parameters were newly introduced, 1.7% to the original (92.9M, without template channels). Notably, to ensure that the model correctly utilizes the pre-trained parameters at the early stage of fine-tuning, we zero-initialized all outputs from the newly introduced modules.

#### Time embedding

UFConf takes a continuous variable *t* ∈ (0, 1] as input. This variable is first encoded with a radial basis function (RBF) and then transformed by a gated linear layer. Specifically, the gated linear layer was defined as ***h*** = sigmoid(***W***_*g*_***x***) ⊙ (***W***_*l*_***x***), where ***W*** s are network parameters and the ⊙ represents element-wise multiplication. These time embedding features are properly reshaped, tiled by numbers of residues and MSAs, and added to the initial MSA and pair representations before inputting them to Evoformer. Notably, the time embedding of residues also indicates missing residues and fixed motifs by setting corresponding representations to *t* = 0 or 1 (see Supplementary A.1).

#### Relative coordinates recycling

In the recycling module, we removed the recycling of MSA and pair representations of AlphaFold2 but retained the recycling of predicted conformations. Specifically, we do not recycle predicted structures 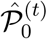, but instead use the module to encode pair-wise distances of the diffused structure 𝒫 ^(*t*)^. We additionally encode relative coordinates between residues, 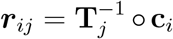, as pair features. These features are processed with RBFs dimension-wise and 2 residual blocks,^39^ and then added to the pair representations as Evoformer inputs. Introducing relative coordinates to Evoformer is crucial because it captures both position and rotation information between residues, providing substantially more information than pair-wise distances alone while keeping all structural inputs to the Evoformer SE(3)-invariant.

#### Structure module

In the original structure module of AlphaFold2, a total of 8 weight-sharing layers transforms a zero-initialized conformation, 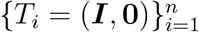, to a predicted conformation 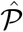. In UFConf, we reuse this module and its pre-trained parameters to iteratively refine 𝒫 ^(*t*)^ into 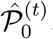. The rationale behind this is that AlphaFold2 uses the learned weights to refine the zero-initialized conformation to the true structure, so we want to best utilize the pre-trained weights to refine noisy structures to multiple possible low-enery structures. Specifically, the diffused positions and rotations were properly scaled and fed to the module. After the alteration, the structural module retains its SE(3)-equivariance, yet a remaining question is whether the pre-trained weights can be smoothly adapted to diffused structures. Surprisingly, our early experiments show the per-trained weights of AlphaFold2 accomplish such robustness even without fine-tuning, as is demonstrated in Supplementary B.4.

### Hierarchical Reweighting of Training Samples

In AlphaFold2 and related protein folding models,^15,20,40^ the sequences of training samples are clustered, and reweighted to make all clusters have uniform weights. This approach has two major drawbacks. First, it is questionable whether uniformly sampling sequence clusters is optimal. For example, this protocol assigns the same weights to the family of antibodies with over 7,000 different sequences, and a rarely studied family with few sequences. This may also be one reason why AlphaFold2 performs unsatisfactorily on local precision of antibody structures.^41^ Second, AlphaFold2 assigns the same sampling weights to conformations within the same sequence cluster, which introduces biases from the uneven data collection practices of the PDB: some conformations are well-studied with abundant structures, while others may have sparser data because they are newly discovered or less studied. Also for homomers, authors may display an assembly either with one chain and roto-translations, or multiple chains of identical conformations, which significantly affects the distribution.

To address the above issues, we developed a hierarchical sampling protocol for UFConf training. We first clustered unique sequences in PDB into sequence clusters 𝒞s representing protein families. We then calculated pairwise TMscores within each cluster and constructed a weighted graph. The Louvain community detection method was then applied to each graph to sub-cluster it into fold clusters ℱs. We finally assign sampling weights hierarchically to each sequence and fold cluster based on the logarithm of their sizes, and each conformation is then reweighted by its size following

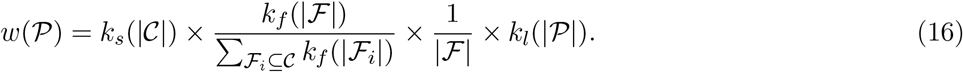

This reweighting protocol balances the distributions of protein families, conformational changes, and lengths in the training data. These balances are especially important for UFConf as a probabilistic generative model, which is expected to diversely sample conformations from the distribution derived from the training data. More details are provided in Supplementary A.2.

### Loss

The general form of loss in UFConf is defined as

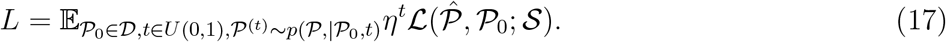

For the choice of ℒ, previous work^12^ shows that using invariant metrics to measure distances between generated and target conformations is inappropriate, as the output conformation should be in the same global frame as the input for correct calculations of score functions. Although the original *frame aligned point error* (FAPE) loss proposed in AlphaFold2 has multiple benefits, it is invariant to roto-translations and thus should be modified. To utilize these benefits while avoiding invariance issues, we add two auxiliary losses with small weights to prevent global shifts: 1) the errors of *C*_*α*_ coordinates between predicted and target conformations (without alignment), and 2) an L2 norm over the structure module’s per-layer updates, which lie in the tangent space at the input structures per-layer. Notably the second loss provides twofold benefits: first, it encourages small changes from the input to avoid unnecessary shifts; second, it regularizes against overfitting and encourages iterative refinements. Other AlphaFold2 losses like pLDDT, distogram and violations are retained. Details are shown in Supplementary A.3.

### Langevin Dynamics and Structural Interpolation

One specific advantage of generating 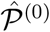 instead of score functions from the model is that this allows new approaches to using the model, such as the generation of continuous trajectories between conformations. We introduce two potential approaches to manipulate UFConf, namely the (overdamped) Langevin dynamics and structural interpolation.

#### Langevin dynamics

The advantage of introducing Langevin dynamics is that this provides a way of recursive, continuous sampling, helpful in studying the dynamic nature of protein systems on larger time scales. Formally, overdamped Langevin dynamics are defined as

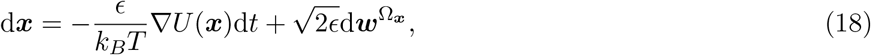

which theoretically converges to a stationary, Boltzmann distribution 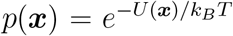. The process can be used to sample from *p*(𝒫, *t*) with *U* (𝒫):= −*k*_*B*_*T* log *p*(𝒫, *t*):

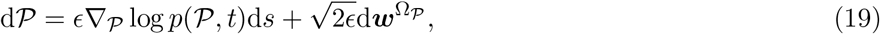

where the scores can be approximated by diffusion models. Notably in Eq. (19), *s* ∈ ℝ is the time of Langevin dynamics, while *t* ∈ (0, 1] is the (fixed) time of diffusion. Prior work^42^ has explored simulating dynamics with learned scores from diffusion models at *t* → 0, but was limited to specific proteins and required massive MD simulations for training. We observed that this framework can be extended to any *t* by revealing the connection between Langevin dynamics and diffusion processes: the Langevin equation is equivalent to moving infinitesimally in the forward diffusion, and then in the reverse direction. This can be immediately proved by observing that Eq. (19) is a direct result of summing Eq. (3) and (4) with *ϵ* = *g*(*t*). More discussions and details are included in Supplementary B.5.

#### Structural interpolation

Another typical application of protein dynamics studies is to find a continuous trajectory of changes from one conformation to another. In computer vision, interpolations of diffused states are commonly used to fuse two images.^32^ In this work, we extend the idea of image interpolation to structural interpolation. For two given conformations 𝒫_0,*A*_ and 𝒫_0,*B*_, the structural interpolation between two structures is done by first diffusing the two backbones to certain time *t* as 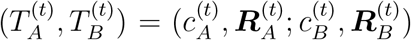, performing linear interpolation in-between as shown in Eq. (20) and Eq. (21), and then integrating the reverse-time diffusion.

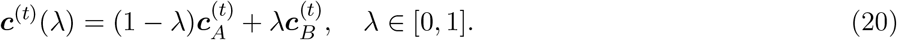

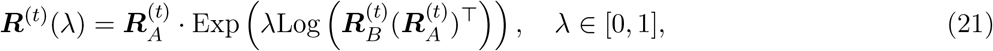

## Data and Software Availability statement

The source codes to run UFConf are available at https://github.com/PKUfjh/Uni-Fold/tree/ufconf, together with the PDB list of the RAC-47 dataset used in the manuscript at https://github.com/PKUfjh/UniFold/blob/ufconf/inference_scripts_ufconf/apo_holo_pairs_info_test47cases.txt. The instruction to run UFConf can be found in https://github.com/PKUfjh/Uni-Fold/blob/ufconf/ufconf/README.md, the checkpoint file to run UFConf can be downloaded in https://zenodo.org/records/11388944.

For other models compared in the manuscript, the source codes to run AlphaFlow are available at https://github.com/bjing2016/alphaflow. The source codes to run AF-cluster are available at https://github.com/HWaymentSteele/AF_Cluster. The source codes to tun MSA-subsampling are available at https://github.com/delalamo/af2_conformations.

The detailed description to generate the RAC-47 dataset are available in Supporting information B.1. The detailed results and visualization of the sampling structures for each case in the RAC-47 dataset are available in Supporting information B.2 and B.6.

## References

(1) Jumper, J.; Evans, R.; Pritzel, A.; Green, T.; Figurnov, M.; Ronneberger, O.; Tunyasuvunakool, K.; Bates, R.; Žídek, A.; Potapenko, A.; others Highly accurate protein structure prediction with AlphaFold. Nature 2021, 596, 583–589.

(2) Oleinikovas, V.; Saladino, G.; Cossins, B. P.; Gervasio, F. L. Understanding cryptic pocket formation in protein targets by enhanced sampling simulations. J. Am. Chem. Soc. 2016, 138, 14257–14263.

(3) Tuckerman, M. E. Statistical mechanics: theory and molecular simulation. 2023,

(4) Sugita, Y.; Okamoto, Y. Replica-exchange molecular dynamics method for protein folding. Chem. Phys. Lett. 1999, 314, 141–151.

(5) Barducci, A.; Bonomi, M.; Parrinello, M. Metadynamics. Wiley Interdisciplinary Reviews: Computational Molecular Science 2011, 1, 826–843.

(6) Kästner, J. Umbrella sampling. Wiley Interdisciplinary Reviews: Computational Molecular Science 2011, 1, 932–942.

(7) Del Alamo, D.; Sala, D.; Mchaourab, H. S.; Meiler, J. Sampling alternative conformational states of transporters and receptors with AlphaFold2. Elife 2022, 11, e75751.

(8) Wayment-Steele, H. K.; Ojoawo, A.; Otten, R.; Apitz, J. M.; Pitsawong, W.; Hömberger, M.; Ovchinnikov, S.; Colwell, L.; Kern, D. Predicting multiple conformations via sequence clustering and AlphaFold2. Nature 2023, 1–3.

(9) Zheng, S.; He, J.; Liu, C.; Shi, Y.; Lu, Z.; Feng, W.; Ju, F.; Wang, J.; Zhu, J.; Min, Y.; others Towards Predicting Equilibrium Distributions for Molecular Systems with Deep Learning. arXiv preprint 2306.05445 2023,

(10) Jing, B.; Berger, B.; Jaakkola, T. AlphaFold Meets Flow Matching for Generating Protein Ensembles. NeurIPS 2023 AI for Science Workshop. 2023.

(11) Abramson, J.; Adler, J.; Dunger, J.; Evans, R.; Green, T.; Pritzel, A.; Ronneberger, O.; Willmore, L.; Ballard, A. J.; Bambrick, J.; others Accurate structure prediction of biomolecular interactions with AlphaFold 3. Nature 2024, 1–3.

(12) Watson, J. L.; Juergens, D.; Bennett, N. R.; Trippe, B. L.; Yim, J.; Eisenach, H. E.; Ahern, W.; Borst, A. J.; Ragotte, R. J.; Milles, L. F.; others De novo design of protein structure and function with RFdiffusion. Nature 2023, 620, 1089–1100.

(13) Yim, J.; Trippe, B. L.; De Bortoli, V.; Mathieu, E.; Doucet, A.; Barzilay, R.; Jaakkola, T. SE (3) diffusion model with application to protein backbone generation. arXiv preprint 2302.02277 2023,

(14) Berman, H. M.; Westbrook, J.; Feng, Z.; Gilliland, G.; Bhat, T. N.; Weissig, H.; Shindyalov, I. N.; Bourne, P. E. The protein data bank. Nucleic acids research 2000, 28, 235–242.

(15) Li, Z.; Liu, X.; Chen, W.; Shen, F.; Bi, H.; Ke, G.; Zhang, L. Uni-Fold: an open-source platform for developing protein folding models beyond AlphaFold. bioRxiv 2022, 2022–08.

(16) Steinegger, M.; Söding, J. MMseqs2 enables sensitive protein sequence searching for the analysis of massive data sets. Nat. Biotechnol. 2017, 35, 1026–1028.

(17) Mirdita, M.; Schütze, K.; Moriwaki, Y.; Heo, L.; Ovchinnikov, S.; Steinegger, M. ColabFold: making protein folding accessible to all. Nat. Methods. 2022, 19, 679–682.

(18) Ying, C.; Cai, T.; Luo, S.; Zheng, S.; Ke, G.; He, D.; Shen, Y.; Liu, T.-Y. Do transformers really perform badly for graph representation? Advances in neural information processing systems 2021, 34, 28877–28888.

(19) Xu, M.; Yu, L.; Song, Y.; Shi, C.; Ermon, S.; Tang, J. Geodiff: A geometric diffusion model for molecular conformation generation. arXiv preprint 2203.02923 2022,

(20) Ahdritz, G.; Bouatta, N.; Kadyan, S.; Xia, Q.; Gerecke, W.; O’Donnell, T. J.; Berenberg, D.; Fisk, I.; Zanichelli, N.; Zhang, B.; others OpenFold: Retraining AlphaFold2 yields new insights into its learning mechanisms and capacity for generalization. bioRxiv 2022, 2022–11.

(21) Boeszoermenyi, A.; Bernaleau, L.; Chen, X.; Kartnig, F.; Xie, M.; Zhang, H.; Zhang, S.; Delacrétaz, M.; Koren, A.; Hopp, A.-K.; others A conformation-locking inhibitor of SLC15A4 with TASL proteostatic anti-inflammatory activity. Nat. Commun. 2023, 14, 6626.

(22) Hassan, N. M.; Alhossary, A. A.; Mu, Y.; Kwoh, C.-K. Protein-ligand blind docking using QuickVina-W with inter-process spatio-temporal integration. Sci. Rep. 2017, 7, 15451.

(23) Koes, D. R.; Baumgartner, M. P.; Camacho, C. J. Lessons learned in empirical scoring with smina from the CSAR 2011 benchmarking exercise. J. Chem. Inf. Model. 2013, 53, 1893–1904.

(24) Corso, G.; Stärk, H.; Jing, B.; Barzilay, R.; Jaakkola, T. Diffdock: Diffusion steps, twists, and turns for molecular docking. arXiv preprint 2210.01776 2022,

(25) Le Guilloux, V.; Schmidtke, P.; Tuffery, P. Fpocket: an open source platform for ligand pocket detection. BMC bioinformatics 2009, 10, 1–11.

(26) Zhou, G.; Gao, Z.; Ding, Q.; Zheng, H.; Xu, H.; Wei, Z.; Zhang, L.; Ke, G. Uni-Mol: A Universal 3D Molecular Representation Learning Framework. The Eleventh International Conference on Learning Representations. 2023.

(27) Alcaide, E.; Li, Z.; Zheng, H.; Gao, Z.; Ke, G. UMD-fit: Generating Realistic Ligand Conformations for Distance-Based Deep Docking Models. NeurIPS 2023 Generative AI and Biology (GenBio) Workshop. 2023.

(28) Müller, C.; Schlauderer, G.; Reinstein, J.; Schulz, G. E. Adenylate kinase motions during catalysis: an energetic counterweight balancing substrate binding. Structure 1996, 4, 147–156.

(29) Evans, R.; O’Neill, M.; Pritzel, A.; Antropova, N.; Senior, A.; Green, T.; Žídek, A.; Bates, R.; Blackwell, S.; Yim, J.; others Protein complex prediction with AlphaFold-Multimer. biorxiv 2021, 2021–10.

(30) Sohl-Dickstein, J.; Weiss, E.; Maheswaranathan, N.; Ganguli, S. Deep unsupervised learning using nonequilibrium thermodynamics. International conference on machine learning. 2015; pp 2256–2265.

(31) Ho, J.; Jain, A.; Abbeel, P. Denoising diffusion probabilistic models. Advances in neural information processing systems 2020, 33, 6840–6851.

(32) Song, Y.; Sohl-Dickstein, J.; Kingma, D. P.; Kumar, A.; Ermon, S.; Poole, B. Score-based generative modeling through stochastic differential equations. arXiv preprint 2011.13456 2020,

(33) Luo, S.; Su, Y.; Peng, X.; Wang, S.; Peng, J.; Ma, J. Antigen-specific antibody design and optimization with diffusion-based generative models for protein structures. Advances in Neural Information Processing Systems 2022, 35, 9754–9767.

(34) Ingraham, J. B.; Baranov, M.; Costello, Z.; Barber, K. W.; Wang, W.; Ismail, A.; Frappier, V.; Lord, D. M.; Ng-Thow-Hing, C.; Van Vlack, E. R.; others Illuminating protein space with a programmable generative model. Nature 2023, 1–9.

(35) Jing, B.; Erives, E.; Pao-Huang, P.; Corso, G.; Berger, B.; Jaakkola, T. EigenFold: Generative Protein Structure Prediction with Diffusion Models. arXiv preprint 2304.02198 2023,

(36) De Bortoli, V.; Mathieu, E.; Hutchinson, M.; Thornton, J.; Teh, Y. W.; Doucet, A. Riemannian score-based generative modelling. Advances in Neural Information Processing Systems 2022, 35, 2406–2422.

(37) Nikolayev, D. I.; Savyolov, T. I.; others Normal distribution on the rotation group SO (3). Texture, Stress, and Microstructure 1997, 29, 201–233.

(38) Nichol, A. Q.; Dhariwal, P. Improved Denoising Diffusion Probabilistic Models. Proceedings of the 38th International Conference on Machine Learning. 2021; pp 8162–8171.

(39) He, K.; Zhang, X.; Ren, S.; Sun, J. Identity Mappings in Deep Residual Networks. Computer Vision – ECCV 2016. Cham, 2016; pp 630–645.

(40) Lin, Z.; Akin, H.; Rao, R.; Hie, B.; Zhu, Z.; Lu, W.; Smetanin, N.; Verkuil, R.; Kabeli, O.; Shmueli, Y.; others Evolutionary-scale prediction of atomic-level protein structure with a language model. Science 2023, 379, 1123–1130.

(41) Polonsky, K.; Pupko, T.; Freund, N. T. Evaluation of the Ability of AlphaFold to Predict the Three-Dimensional Structures of Antibodies and Epitopes. J. Immun. 2023, 211, 1578–1588.

(42) Arts, M.; Garcia Satorras, V.; Huang, C.-W.; Zügner, D.; Federici, M.; Clementi, C.; Noé, F.; Pinsler, R.; van den Berg, R. Two for one: Diffusion models and force fields for coarse-grained molecular dynamics. J. Chem. Theory Comput. 2023, 19, 6151–6159.

