## Supplementary information file for "Accurate Conformation Sampling via Protein Structural Diffusion"

### A Implementation Details

#### A.1 Markovian Span-Masking and Missing Residues

To recap, the span-masking strategy is defined as:

$$M_i \sim \text{Bernoulli}(p_i),\ i \geq 1, \qquad p_1 = \rho,$$
$$p_i = \frac{K-1}{K} M_{i-1} + \frac{\rho}{(1-\rho)K}(1 - M_{i-1}),\ i > 1,$$

where $M_i = 1$ indicates residue $i$ is selected as a motif and not diffused during the forward process. Notably, when $i$ is a residue without experimentally solved structure, $M_i$ is always set as 0.

We now prove that 1) the overall masking rate is $\rho$, i.e., $\mathbb{E}(\frac{1}{n}\sum_{i=1}^{n} M_i) = \rho$ and 2) the expected number of consecutive motifs is $K$. To show 1), by induction it can be easily verified that $P(M_i = 1) = \rho$. Therefore,

$$\mathbb{E}\left(\frac{1}{n}\sum_{i=1}^{n} M_i\right) = \frac{1}{n}\sum_{i=1}^{n} P(M_i = 1) = \rho. \tag{S1}$$

For 2), note that $P(M_{i+1} = 1|M_i = 1) = 1 - 1/K$. Therefore, conditioned on $M_i = 1$, the length of the sequence of consecutive 1s $(i, i+1, \cdots)$ follows a Geometric($\frac{1}{K}$) distribution, and the expectation can be calculated as $K$.

In the training of UFConf, we set $K = 15$ and calculated $\rho$ per sample as

$$\rho = \max(0, \tilde{\rho} - (1 - r)), \quad \tilde{\rho} \sim \text{Uniform}(0, 1), \quad r = 0.6. \tag{S2}$$

Additionally, no extra symbols were introduced to encode motifs. Instead, we assigned $t_i = 0$ to fully leverage the time embedding to represent that the residues have zero noise. We took a similar approach for missing residues without solved structures, drawing their structures from the prior and assigning $t_i = 1$.

#### A.2 Hierarchical Reweighting Details

We collected all protein chains that were released before April 30, 2022, and filtered out those without valid sequences. A total of 645,947 protein chains, with 133,954 unique sequences were collected. To cluster these unique sequences, we used MMseqs2[1] with parameters `-min-seq-id 0.4 -c 0.8 -cov-mode 1`, which created sequence clusters sharing $> 40\%$ sequence identities. This generated 37,838 sequence clusters.

For the 22,122 sequence clusters with at least 3 sequences, we further divided them into fold clusters by graph partition. We used TMalign[2] to calculate all pairwise TMscores inside each sequence cluster based on the implementation of Foldseek.[3] Notably, this implementation prefilters and avoids calculating TMscores for pairs unlikely to have high similarity. We assigned a TMscore of 0 for such pairs. All calculated TMscores $(r)$ were processed into link weights via the kernel

$$w = \left(\frac{\max(0, r - r_0)}{1 - r_0}\right)^m \tag{S3}$$

with $r_0 = 0.85$ and $m = 2$. Notably, this kernel ignores connections between structures with $r < 0.85$. We partitioned the generated graph using the Louvain algorithm[4] to produce communities, i.e., the fold clusters. This generated 41,567 fold clusters.

To recap, the final sampling weight $w(\mathcal{P})$ of each conformation $\mathcal{P} \in \mathcal{F}, \mathcal{F} \subseteq \mathcal{C}$ was calculated as:

$$w(\mathcal{P}) = k_s(|\mathcal{C}|) \times \frac{k_f(|\mathcal{F}|)}{\sum_{\mathcal{F}_i \subseteq \mathcal{C}} k_f(|\mathcal{F}_i|)} \times \frac{1}{|\mathcal{F}|} \times k_l(|\mathcal{P}|).$$

Specifically, we used:

$$k_s(x) = \log_{10}(x + 9) \tag{S4}$$

$$k_f(x) = \log_{10}(x + 99) \tag{S5}$$

$$k_l(x) = \frac{1}{2}\left(1 - \cos\left(\frac{\pi}{2} \cdot \max(512, \min(x, 32))\right)\right) \tag{S6}$$

Intuitively, the sampling probability of each cluster is still relative to its size, but much smoothed by logarithm. The bias to $k_f$ is larger than $k_s$ because we want the conformations to be sampled more evenly (weights ranging from 2.0 to 3.5). Sequences with very short lengths are punished by $k_l$, keeping a minimal sampling weight $\approx 0.01$. Fig. S1 shows more details on the distributions and impacts of so-defined sampling protocol.

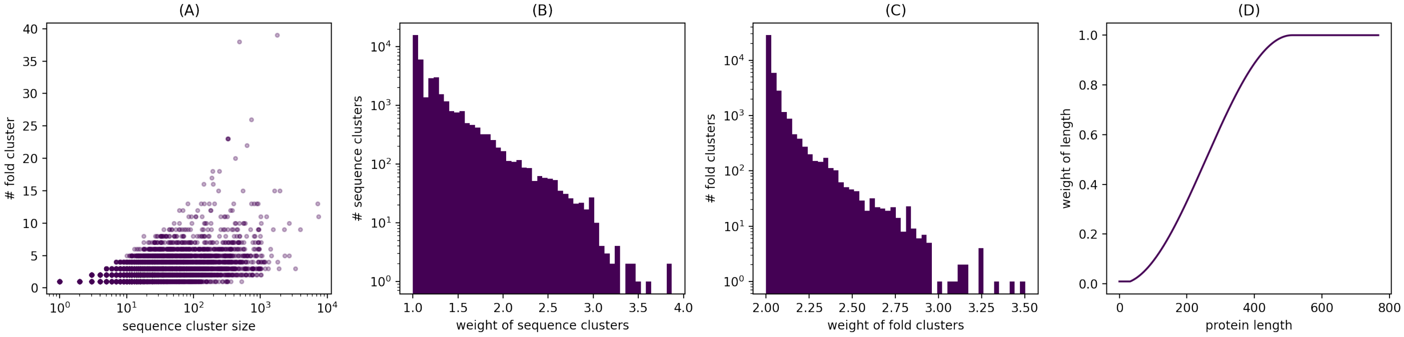


Figure S1: Details of the hierarchical reweighting protocol. (A) number of fold clusters versus sequence cluster sizes. (B) distribution of sequence cluster weights; (C) distribution of fold cluster weights; (D) length weight versus protein length.

#### A.3 Loss

As is discussed in Section 4.5, we use the original losses introduced in AlphaFold as well as 2 additional losses, namely a loss of $C_\alpha$ coordinate errors and a penalty over the update vector. The formal definition of the loss we use is

$$\mathcal{L} = \mathcal{L}_{\text{FAPE}} + \mathcal{L}_{\text{chi}} + 0.1\mathcal{L}_{\text{penalty}} + 0.1\mathcal{L}_{\text{ca-error}} + 0.02\mathcal{L}_{\text{violation}} + \mathcal{L}_{\text{aux}}, \tag{S7}$$

$$\mathcal{L}_{\text{aux}} = 2.0\mathcal{L}_{\text{BERT-MSA}} + 0.3\mathcal{L}_{\text{distogram}} + 0.01\mathcal{L}_{\text{plddt}} + 0.1\mathcal{L}_{\text{pae}} + 0.01\mathcal{L}_{\text{resolved}}, \tag{S8}$$

in which we inherited most of the loss weights in AlphaFold's pre-training. The definition of $\mathcal{L}_{\text{penalty}}$ and $\mathcal{L}_{\text{ca-error}}$ are

$$\mathcal{L}_{\text{ca-error}} = \frac{1}{n}\sum_{i=1}^{n} \|\hat{\boldsymbol{c}}_i - \boldsymbol{c}_i\|_2 \tag{S9}$$

$$\mathcal{L}_{\text{penalty}} = \frac{1}{n}\sum_{i=1}^{n}\sum_{l=1}^{L} \left(\|\hat{\boldsymbol{c}}_i^{(l)} - \hat{\boldsymbol{c}}_i^{(l-1)}\|_2 + \|\boldsymbol{q}_i^{(l)}\|_2\right), \qquad \hat{\boldsymbol{c}}_i^{(l)} = \boldsymbol{c}^{(t)} \tag{S10}$$

where $\boldsymbol{q}_i \in \mathbb{R}^3$ is the vector of (unnormalized) imaginary components in the quaternion representation of the update rotation. Notably, $\mathcal{L}_{\text{FAPE}}, \mathcal{L}_{\text{chi}}, \mathcal{L}_{\text{penalty}}$ and $\mathcal{L}_{\text{ca-error}}$ are calculated for per-layer outputs of the structure module. The FAPE loss of both backbones and sidechains are all calculated and weighted as $\mathcal{L}_{\text{FAPE}} = 0.5(\mathcal{L}_{\text{bb}} + \mathcal{L}_{\text{sc}})$.

#### A.4 Training and Inference Algorithm

We formulate the training algorithm of UFConf as Algorithm 1. For inference, we first introduce the discretization of the continuous form of the reverse-time SDE. We implemented the specific discretization form as

$$\boldsymbol{c}^{(t-\tau)} = \frac{1}{\sqrt{1-\beta_{\text{pos}}^{(t,t-\tau)}}}\left(\boldsymbol{c}^{(t)} + \beta_{\text{pos}}^{(t,t-\tau)}\nabla_{\boldsymbol{c}^{(t)}}\log p(\boldsymbol{c}^{(t)}, t|\hat{\boldsymbol{c}}^{(t)})\right) + \frac{1-\gamma_{\text{pos}}^{(t-\tau)}}{(1-\gamma_{\text{pos}}^{(t)})\beta_{\text{pos}}^{(t,t-\tau)}}\boldsymbol{\epsilon}_{\text{pos}}, \quad \boldsymbol{\epsilon}_{\text{pos}} \sim \mathcal{N}(\boldsymbol{0}, \boldsymbol{I}_3) \tag{S11}$$

$$\boldsymbol{R}^{(t-\tau)} = \text{Exp}\left(\beta_{\text{rot}}^{(t,t-\tau)}\nabla_{\boldsymbol{R}^{(t)}}\log p(\boldsymbol{R}^{(t)}, t|\hat{\boldsymbol{R}}^{(t)}) + \boldsymbol{\epsilon}_{\text{rot}}^{(t)}\right)\cdot\boldsymbol{R}^{(t)}, \quad \boldsymbol{\epsilon}_{\text{rot}}^{(t)} \sim \text{IGSO3}\left(\beta_{\text{rot}}^{(t,t-\tau)}\right). \tag{S12}$$

Specifically, Eq. (S11) is a reparameterization of the classic DDPM[5] instead of VP-SDE,[6] because we find that the discretization form of VP-SDE suffers from numerical issues when $\beta \gg 0$. When drawing $\boldsymbol{\epsilon}_{\text{rot}} \sim \text{IGSO3}(\beta)$, we sample uniformly distributed rotation axis from the unit ball of $\mathbb{R}^3$ and sample a rotation angle from $\psi(\cdot,\beta)$.The parameters of $\beta_*^{(t,t-\tau)}$ are calculated as

$$\beta_{\text{pos}}^{(t,t-\tau)} = 1 - \frac{\gamma_{\text{pos}}^{(t)}}{\gamma_{\text{pos}}^{(t-\tau)}}, \tag{S13}$$

$$\beta_{\text{rot}}^{(t,t-\tau)} = \sigma_{\text{rot}}^{(t)} - \sigma_{\text{rot}}^{(t-\tau)}, \tag{S14}$$

which corresponds to the definitions of $\gamma$ and $\sigma$s. The detailed algorithm of inference is presented in Algorithm 3.

---

**Algorithm 1** UFConf training.

---

**Input:** dataset $\mathcal{D} = \{(\mathcal{S},\mathcal{P})\}$, pre-trained AlphaFold weights $\theta$
Initialize model weights as $\theta$.
**repeat**
  Sample $(\mathcal{S},\mathcal{P})$ from $\mathcal{D}$ according to Eq.(16).
  Sample $t \sim \text{Uniform}(0,1)$.
  Sample motif mask $\mathcal{M}$ according to Eq.(15).
  $T^{(t)} = \text{ForwardDiffusion}(t,\mathcal{P},\mathcal{M})$ (see Algorithm 2).
  $\hat{\mathcal{P}} = m_\theta(t, T^{(t)}, \mathcal{S})$.
  Calculate $\mathcal{L}(\hat{\mathcal{P}},\mathcal{P};\mathcal{S})$ following Eq. (S7).
  Backpropagate $\mathcal{L}$ to update $\theta$.
**until** convergence

---

**Algorithm 2** ForwardDiffusion

---

**Input:** time $t \in (0,1]$, protein $\mathcal{P} = (\mathcal{A}_i = ((\boldsymbol{R}_i, \boldsymbol{c}_i), \mathcal{X}_i))_{i=1}^n$, motif mask $\mathcal{M} = (M_1, \cdots, M_n)$
**for** $i = 1$ **to** $n$ **do**
  **if** $M_i = 0$ (residue not fixed as motif) **then**
    **if** structure of $\mathcal{A}_i$ is missing **then**
      Draw $T_i^{(t)} = (\boldsymbol{R}_i^{(t)}, \boldsymbol{c}_i^{(t)})$ from prior.
    **else**
      Sample $T_i^{(t)} = ((\boldsymbol{R}_i^{(t)}, \boldsymbol{c}_i^{(t)}))$ following Eq. (7)(8) providing $\boldsymbol{R}_i, \boldsymbol{c}_i$.
    **end if**
  **else**
    $T_i^{(t)} = T_i = (\boldsymbol{R}_i, \boldsymbol{c}_i)$
  **end if**
**end for**
**Return:** $T^{(t)} = \left\{T_i^{(t)}\right\}_{i=1}^n$

---

**Algorithm 3** UFConf inference.

---

**Input:** protein sequence $\mathcal{S}$, motif mask $\mathcal{M}$, motif structures $\{T_i^{\text{M}} | M_i = 1\}$, number of steps $S$
$t = 1, \tau = 1/S$.
Initialize $T^{(1)}$ given $\mathcal{M}, \{T_i^{\text{M}} | M_i = 1\}$ (see Algorithm 4).
**for** $s = 1$ **to** $S$ **do**
  Calculate $\hat{\mathcal{P}} = m_\theta(t, T^{(t)}, \mathcal{S})$.
  $T^{(t-\tau)} = \text{ReverseDiffusion}(t, \tau, T^{(t)}, \hat{\mathcal{P}}, \mathcal{M})$ (see Algorithm 5).
  $t = t - \tau$
**end for**
**Return** $\mathcal{P}^{(0)}$

---

**Algorithm 4** Initialize

**Input:** motif mask $\mathcal{M} = (M_1, \cdots, M_n)$, motif structures $\{\mathcal{A}_i^{\mathrm{M}} | M_i = 1\}$
**for** $i = 1$ **to** $n$ **do**
  **if** $M_i = 0$ (not provided as motif) **then**
    Draw $T_i^{(t)} = (\boldsymbol{R}_i^{(t)}, \boldsymbol{c}_i^{(t)})$ from prior.
  **else**
    $T_i = T_i^{\mathrm{M}}$
  **end if**
**end for**
**Return** $T^{(1)} = (T_1, \cdots, T_n)$

**Algorithm 5** ReverseDiffusion

**Input:** time $t \in (0, 1]$, interval $\tau \in (0, t]$, diffused protein backbone $T^{(t)}$, predicted protein $\hat{\mathcal{P}}$, motif mask $\mathcal{M}$.
**if** $t - \tau = 0$ **then**
  $\mathcal{P}^{(t-\tau)} = \hat{\mathcal{P}}$
**else**
  **for** $i = 1$ **to** $n$ **do**
    **if** $M_i = 0$ (not provided as motif) **then**
      Calculate $T_i^{(t-\tau)}$ with Eq. (S11) - (S12) providing $T_i^{(t)}, \hat{T}_i, t, \tau$.
    **else**
      $T_i^{(t-\tau)} = T_i^{(t)}$
    **end if**
  **end for**
**end if**
**Return** $T^{(t-\tau)} = (T_1^{(t-\tau)}, \cdots, T_n^{(t-\tau)})$

#### A.5 Training Details

The structural diffusion fine-tuning was initialized with parameters in Model 2 of the v3 parameters of AlphaFold-Multimer [1]. Table S1 summarizes the parameters used to train UFConf. Most of the settings are inherited from AlphaFold-Multimer (v3). The original parameters were adapted to Uni-Fold[7] implementation[2] using the provided utilities in the repository. Training data and generated MSAs were downloaded from the repository as well. Due to the cost of fine-tuning, we did not do a thorough hyperparameter search.

### B More Results

#### B.1 RAC-47 Curation Details

To curate RAC-47, we collected all 82777 protein chains in PDB[8] released between 2022-04-30 and 2024-03-29. MMseqs2[1] was used to cluster the sequences with 100% sequence identity and clusters with sizes ranging from 2 to 10 were kept, resulting in 28409 chains. We then filtered for chains with 128 to 768

[1] https://storage.googleapis.com/alphafold/alphafold_params_2022-12-06.tar
[2] https://github.com/dptech-corp/Uni-Fold/

Table S1: Details of parameters in UFConf.

| Parameters | Values |
|---|---|
| number of samples | 645,947 |
| batch size | 64 |
| total steps | 20,000 |
| peak learning rate | 1e-3 |
| warm-up steps | 1000 |
| learning rate decay | N/A |
| number of MSAs | 512 |
| number of extra MSAs | 2,048 |
| sequence crop size | 256 |
| number of templates | N/A |
| dimension of MSA, pair and single representations | (256, 128, 384) |
| dimension of RBF of time | 512 |
| dimension of RBF of relative coordinates | 64 |

residues, yielding a set of 20078 chains. From the initially filtered 20078 protein structures, we identified 4256 sequence clusters, each containing at least one of these structures. Aligned backbone Root Mean Square Deviations (RMSDs) for all pairings within these clusters were calculated using PyMOL.[9] The primary selection criterion targeted clusters with at least one pair of RMSD greater than 2 Angstroms. This process resulted in 304 clusters of chains. But among these clusters, there exists clusters where the resolved structures have rather different sequences (due to the experimental details). For convenience we just select those clusters with the resolved structures having the same sequence, this procedure produces 55 clusters. Finally, we exclude the clusters where the resolved structures have incontinuous sequence, our pipeline yielded 47 unbiased protein clusters for analysis.

#### B.2 Detailed results and discussions on RAC-47

In Table S2 we show the best TM-score among 100 generated conformations corresponding to two real conformations with the largest RMSD for each case in our **RAC-47** dataset. Note that some cases are neglected for AF-cluster since there are no clusters generated using the default settings in AF-cluster.

Although the RAC-47 dataset is created by selecting the sequence clusters outside the training dataset, there may be doubts that the training dataset may contain protein sequences with high similarities (>90%) to those in the RAC-47 dataset. If so, then the model is argued to be just remembering the structure information in the training dataset rather than generalizing to the whole new sequences.

Table S2: Evaluation results on RAC-47. *Diff.* indicates the TM-score between two real conformations. The best TM-score of 100 generated conformations to both of the real ones in each case are reported and aggregated. MSA-sub. represents MSA-subsampling method.

| PDB IDs | Diff | AF2 | AF3 | UFConf | AlphaFlow | MSA-sub. (64) | MSA-sub. (256) | AF-cluster |
|---|---|---|---|---|---|---|---|---|
| 7Z3N_C/7Z3O_C | 0.658 | 0.567/0.651 | 0.535/0.670 | 0.634/0.571 | 0.558/0.673 | 0.556/0.638 | 0.566/0.634 | 0.498/0.598 |
| 7R5J_F0/7R5K_F0 | 0.461 | 0.388/0.387 | 0.405/0.417 | 0.395/0.392 | 0.418/0.428 | 0.389/0.407 | 0.395/0.390 | 0.354/0.392 |
| 7YCO_B/8H6F_X | 0.543 | 0.966/0.538 | 0.966/0.541 | 0.968/0.556 | 0.971/0.547 | 0.973/0.550 | 0.970/0.539 | 0.955/0.576 |
| 8TOC_b/8TV9_a | 0.658 | 0.411/0.402 | 0.450/0.473 | 0.392/0.396 | 0.386/0.418 | 0.433/0.424 | 0.408/0.408 | N/A |
| 8TH8_S/8TID_s | 0.527 | 0.537/0.551 | 0.524/0.557 | 0.621/0.659 | 0.603/0.690 | 0.544/0.563 | 0.559/0.576 | 0.561/0.576 |
| 7P37_A/7P3F_A | 0.657 | 0.726/0.808 | 0.749/0.828 | 0.784/0.823 | 0.727/0.821 | 0.753/0.706 | 0.731/0.778 | 0.731/0.725 |
| 7UTI_Z/7UTL_d | 0.561 | 0.505/0.446 | 0.502/0.449 | 0.483/0.592 | 0.574/0.551 | 0.508/0.445 | 0.499/0.460 | 0.512/0.512 |
| 8BDV_A/8BH7_A | 0.537 | 0.422/0.596 | 0.642/0.647 | 0.573/0.771 | 0.620/0.787 | 0.443/0.675 | 0.502/0.732 | 0.663/0.804 |
| 7SJO_H/7SJP_H | 0.651 | 0.875/0.879 | 0.949/0.842 | 0.915/0.906 | 0.946/0.932 | 0.841/0.923 | 0.841/0.882 | 0.815/0.883 |
| 7TUC_A/7TUE_A | 0.716 | 0.981/0.702 | 0.947/0.696 | 0.953/0.677 | 0.990/0.739 | 0.979/0.758 | 0.978/0.717 | 0.929/0.737 |
| 8DKF_H/8DOW_C | 0.679 | 0.903/0.770 | 0.932/0.932 | 0.905/0.911 | 0.945/0.919 | 0.896/0.818 | 0.903/0.805 | 0.679/0.688 |
| 8EFR_A/8EFT_A | 0.787 | 0.842/0.759 | 0.851/0.762 | 0.812/0.795 | 0.836/0.802 | 0.836/0.738 | 0.856/0.772 | 0.405/0.404 |
| 8E2Y_A/8E31_A | 0.674 | 0.727/0.648 | 0.742/0.654 | 0.752/0.669 | 0.752/0.663 | 0.705/0.635 | 0.730/0.655 | 0.629/0.578 |
| 8D01_L/8D0Y_L | 0.698 | 0.876/0.870 | 0.953/0.906 | 0.950/0.919 | 0.912/0.876 | 0.967/0.723 | 0.930/0.863 | 0.835/0.780 |
| 8BFL_A/8BFP_A | 0.739 | 0.892/0.860 | 0.947/0.831 | 0.929/0.911 | 0.855/0.895 | 0.874/0.878 | 0.880/0.910 | 0.271/0.280 |
| 7SJO_F/7SJP_L | 0.699 | 0.925/0.956 | 0.925/0.982 | 0.942/0.983 | 0.956/0.830 | 0.880/0.973 | 0.923/0.970 | 0.621/0.743 |
| 8DUE_A/8DVF_A | 0.720 | 0.775/0.790 | 0.833/0.894 | 0.822/0.944 | 0.882/0.934 | 0.844/0.801 | 0.810/0.807 | 0.449/0.459 |
| 7ZWM_E/7ZXF_E | 0.712 | 0.937/0.958 | 0.911/0.982 | 0.841/0.980 | 0.925/0.984 | 0.830/0.976 | 0.947/0.977 | 0.623/0.707 |
| 7Y5A_D/7Y5B_D | 0.829 | 0.893/0.935 | 0.899/0.930 | 0.943/0.961 | 0.945/0.948 | 0.916/0.940 | 0.894/0.939 | 0.847/0.923 |
| 8FWF_L/8FYM_L | 0.749 | 0.957/0.919 | 0.968/0.883 | 0.981/0.896 | 0.985/0.978 | 0.983/0.949 | 0.974/0.925 | 0.722/0.623 |
| 8B6V_A/8B6W_A | 0.655 | 0.964/0.669 | 0.925/0.626 | 0.973/0.707 | 0.956/0.750 | 0.950/0.758 | 0.969/0.706 | 0.362/0.359 |
| 8DKF_L/8DOW_D | 0.764 | 0.943/0.955 | 0.982/0.952 | 0.981/0.962 | 0.972/0.957 | 0.979/0.943 | 0.984/0.957 | 0.916/0.889 |
| 7WKP_A/7WWU_I | 0.723 | 0.921/0.695 | 0.925/0.701 | 0.950/0.730 | 0.926/0.724 | 0.929/0.705 | 0.937/0.699 | 0.925/0.689 |
| 7PIS_9/7PIT_9 | 0.872 | 0.721/0.716 | 0.889/0.910 | 0.868/0.916 | 0.885/0.909 | 0.878/0.921 | 0.871/0.911 | 0.290/0.297 |
| 8DKE_P/8DKI_P | 0.813 | 0.963/0.963 | 0.862/0.969 | 0.949/0.941 | 0.953/0.960 | 0.963/0.945 | 0.947/0.968 | 0.960/0.855 |
| 8G4C_D/8G4D_D | 0.808 | 0.856/0.841 | 0.887/0.859 | 0.916/0.883 | 0.879/0.847 | 0.845/0.787 | 0.864/0.829 | 0.672/0.630 |
| 7ZF5_D/7ZF6_H | 0.801 | 0.956/0.937 | 0.977/0.923 | 0.924/0.944 | 0.972/0.901 | 0.928/0.933 | 0.910/0.950 | 0.676/0.620 |
| 8HKX_AS7P/8HKY_AS7P | 0.755 | 0.797/0.771 | 0.797/0.775 | 0.803/0.796 | 0.800/0.788 | 0.782/0.748 | 0.790/0.763 | 0.782/0.755 |
| 8HFX_E/8IFY_E | 0.890 | 0.915/0.888 | 0.917/0.892 | 0.927/0.903 | 0.916/0.886 | 0.918/0.893 | 0.921/0.893 | 0.905/0.877 |
| 8SQZ_C/8SRM_C | 0.754 | 0.687/0.862 | 0.676/0.856 | 0.684/0.846 | 0.709/0.859 | 0.691/0.862 | 0.689/0.864 | 0.682/0.855 |
| 8HKX_S13P/8HKY_S13P | 0.769 | 0.786/0.768 | 0.795/0.785 | 0.797/0.772 | 0.801/0.766 | 0.788/0.759 | 0.782/0.770 | 0.741/0.731 |
| 8HKY_AL1P/8HKZ_AL1P | 0.826 | 0.794/0.770 | 0.801/0.778 | 0.816/0.768 | 0.825/0.784 | 0.793/0.776 | 0.800/0.777 | 0.795/0.776 |
| 8V2D_a/8V3B_A | 0.874 | 0.826/0.860 | 0.851/0.899 | 0.817/0.813 | 0.944/0.907 | 0.835/0.863 | 0.829/0.861 | N/A |
| 8E39_A/8E3A_A | 0.757 | 0.724/0.649 | 0.728/0.660 | 0.733/0.683 | 0.783/0.696 | 0.672/0.670 | 0.735/0.681 | 0.533/0.507 |
| 8TCA_L/8VEV_B | 0.840 | 0.895/0.978 | 0.790/0.954 | 0.951/0.976 | 0.960/0.967 | 0.975/0.971 | 0.906/0.976 | 0.899/0.932 |
| 8EE5_L/8EF3_B | 0.843 | 0.696/0.871 | 0.827/0.699 | 0.807/0.954 | 0.952/0.980 | 0.912/0.959 | 0.929/0.933 | 0.807/0.940 |
| 8EE5_H/8EF3_A | 0.846 | 0.824/0.681 | 0.928/0.865 | 0.936/0.908 | 0.966/0.956 | 0.866/0.812 | 0.844/0.721 | 0.838/0.778 |
| 7ZF5_F/7ZF6_L | 0.856 | 0.976/0.949 | 0.973/0.990 | 0.976/0.992 | 0.990/0.959 | 0.967/0.980 | 0.989/0.970 | 0.665/0.701 |
| 8HC3_E/8HC6_L | 0.853 | 0.898/0.889 | 0.914/0.905 | 0.916/0.917 | 0.934/0.932 | 0.898/0.905 | 0.856/0.827 | 0.893/0.865 |
| 7QD7_AAA/7QH9_AAA | 0.900 | 0.985/0.957 | 0.982/0.964 | 0.985/0.981 | 0.984/0.973 | 0.977/0.960 | 0.983/0.954 | 0.305/0.306 |
| 7T22_A/7T23_B | 0.707 | 0.849/0.707 | 0.860/0.702 | 0.931/0.870 | 0.879/0.753 | 0.821/0.811 | 0.836/0.704 | 0.804/0.722 |
| 8UOP_A/8UOY_B | 0.898 | 0.907/0.936 | 0.916/0.944 | 0.918/0.943 | 0.918/0.941 | 0.921/0.936 | 0.912/0.936 | 0.907/0.934 |
| 7S8G_L/7UDS_L | 0.856 | 0.866/0.923 | 0.879/0.937 | 0.935/0.947 | 0.975/0.962 | 0.736/0.676 | 0.773/0.700 | 0.827/0.895 |
| 8I6O_B/8I6Q_B | 0.862 | 0.895/0.916 | 0.906/0.920 | 0.943/0.925 | 0.928/0.920 | 0.908/0.918 | 0.908/0.915 | 0.915/0.916 |
| 8GAE_E/8GFT_E | 0.898 | 0.674/0.675 | 0.675/0.676 | 0.676/0.678 | 0.676/0.675 | 0.674/0.675 | 0.674/0.675 | 0.672/0.675 |
| 8HC3_H/8HC5_H | 0.878 | 0.868/0.840 | 0.892/0.933 | 0.909/0.912 | 0.875/0.894 | 0.819/0.775 | 0.856/0.865 | 0.634/0.668 |
| 8JSG_A/8JSH_A | 0.849 | 0.816/0.730 | 0.876/0.815 | 0.820/0.762 | 0.828/0.761 | 0.863/0.809 | 0.843/0.773 | 0.808/0.798 |
| Mean-of-Best | 0.753 | 0.811/0.783 | 0.831/0.803 | 0.839/0.824 | 0.850/0.826 | 0.818/0.793 | 0.822/0.794 | 0.696/0.688 |
| Median-of-Best | 0.757 | 0.866/0.808 | 0.887/0.856 | 0.915/0.896 | 0.916/0.876 | 0.863/0.809 | 0.856/0.807 | 0.722/0.722 |

To remove this possibility, we identified all sequence clusters containing the RAC-47 dataset with 90% identity, and then removed the clusters which contain chains in the training dataset. After this process we get 30 sequence clusters termed as **RAC-30**, by construction the training dataset does not contain any sequences with high similarities (>90%) to those in this new benchmark dataset.

Since the RAC-30 dataset is the subset of the RAC-47, we can simply extract the results for this RAC-30 dataset from the original RAC-47 dataset for each model. As shown in Table S3, the results in RAC-30 have the same implication with RAC-47: UFConf performs the best in terms of the median TM-score, while AlphaFlow achieves the best mean TM-score, and UFConf achieves the most successful sampling with 15 among 30 cases (which shows an increase of the success ratio w.r.t RAC-47). Regarding recovery, UFConf and AlphaFlow performs the best in terms of median and mean MAT-R respectively. In Table S4 we show the best TM-score among 100 generated conformations corresponding to two real conformations with the largest RMSD for each case in the RAC-30 dataset.

Table S3: Evaluation results on RAC-30. Mean *Diff.* indicates the mean TM-score between two real conformations across 30 cases. The best TM-score of 100 generated conformations to both of the real ones in each case are reported and aggregated. The depth of MSA for MSA-subsampling model is indicated in the bracket.

| - | Mean *Diff.* | Alphafold2 | Alphafold3 | UFConf | AlphaFlow |
|---|---|---|---|---|---|
| Mean-of-Best TM-score (↑) | 0.742 | 0.796/0.784 | 0.825/0.817 | 0.832/0.829 | **0.837**/**0.834** |
| Median-of-Best TM-score (↑) | 0.754 | 0.861/0.824 | 0.888/0.862 | **0.912**/**0.907** | 0.899/0.885 |
| Num. of successful sampling (↑) | - | 5 | 8 | **15** | 11 |
| mean/median MAT-R (↑) | - | 0.794/0.824 | 0.827/0.879 | 0.836/**0.908** | **0.838**/0.876 |
| mean/median MAT-P (↑) | - | 0.788/0.839 | 0.809/0.870 | 0.739/0.789 | 0.791/0.832 |

| - | MSA-subsampling (64) | MSA-subsampling (256) | AF-cluster |
|---|---|---|---|
| Mean-of-Best TM-score (↑) | 0.801/0.785 | 0.803/0.793 | 0.654/0.659 |
| Median-of-Best TM-score (↑) | 0.845/0.794 | 0.856/0.817 | 0.677/0.707 |
| Num. of successful sampling (↑) | 5 | 6 | 1 |
| mean/median MAT-R (↑) | 0.798/0.839 | 0.802/0.837 | 0.660/0.711 |
| mean/median MAT-P (↑) | 0.736/0.769 | 0.778/0.810 | 0.469/0.502 |

#### B.3 Results of existing docking methods in redocking setup

The results of three docking methods are shown in Figure Fig. S2, the binding ligand RMSD with the experimental structure is 5.997 Å for QuickVina-W, 6.633 Å for Smina and 8.543 Å for Diffdock. The results indicate that the three docking methods fail to dock the ligand to the right binding mode using the experimental structure (8JZX).

Table S4: Evaluation results on RAC-30. *Diff.* indicates the TM-score between two real conformations. The best TM-score of 100 generated conformations to both of the real ones in each case are reported and aggregated. MSA-sub. represents MSA-subsampling method.

| PDB IDs | Diff | AF2 | AF3 | UFConf | AlphaFlow | MSA-sub. (64) | MSA-sub. (256) | AF-cluster |
|---|---|---|---|---|---|---|---|---|
| 7Z3N_C/7Z3O_C | 0.658 | 0.567/0.651 | 0.535/0.670 | 0.634/0.571 | 0.558/0.673 | 0.556/0.638 | 0.566/0.634 | 0.498/0.598 |
| 7R5J_F0/7R5K_F0 | 0.461 | 0.388/0.387 | 0.405/0.417 | 0.395/0.392 | 0.418/0.428 | 0.389/0.407 | 0.395/0.390 | 0.354/0.392 |
| 8TOC_b/8TV9_a | 0.658 | 0.411/0.402 | 0.450/0.473 | 0.392/0.396 | 0.386/0.418 | 0.433/0.424 | 0.408/0.408 | N/A |
| 8TH8_S/8TID_s | 0.527 | 0.537/0.551 | 0.524/0.557 | 0.621/0.659 | 0.603/0.690 | 0.544/0.563 | 0.559/0.576 | 0.561/0.576 |
| 7P37_A/7P3F_A | 0.657 | 0.726/0.808 | 0.749/0.828 | 0.784/0.823 | 0.727/0.821 | 0.753/0.706 | 0.731/0.778 | 0.731/0.725 |
| 8BDV_A/8BH7_A | 0.537 | 0.422/0.596 | 0.642/0.647 | 0.573/0.771 | 0.620/0.787 | 0.443/0.675 | 0.502/0.732 | 0.663/0.804 |
| 7SJO_H/7SJP_H | 0.651 | 0.875/0.879 | 0.949/0.842 | 0.915/0.906 | 0.946/0.932 | 0.841/0.923 | 0.841/0.882 | 0.815/0.883 |
| 8DKF_H/8DOW_C | 0.679 | 0.903/0.770 | 0.932/0.932 | 0.905/0.911 | 0.945/0.919 | 0.896/0.818 | 0.903/0.805 | 0.679/0.688 |
| 8EFR_A/8EFT_A | 0.787 | 0.842/0.759 | 0.851/0.762 | 0.812/0.795 | 0.836/0.802 | 0.836/0.738 | 0.856/0.772 | 0.405/0.404 |
| 8D01_L/8D0Y_L | 0.698 | 0.876/0.870 | 0.953/0.906 | 0.950/0.919 | 0.912/0.876 | 0.967/0.723 | 0.930/0.863 | 0.835/0.780 |
| 8BFL_A/8BFP_A | 0.739 | 0.892/0.860 | 0.947/0.831 | 0.929/0.911 | 0.855/0.895 | 0.874/0.878 | 0.880/0.910 | 0.271/0.280 |
| 7SJO_F/7SJP_L | 0.699 | 0.925/0.956 | 0.925/0.982 | 0.942/0.983 | 0.956/0.830 | 0.880/0.973 | 0.923/0.970 | 0.621/0.743 |
| 8DUE_A/8DVF_A | 0.720 | 0.775/0.790 | 0.833/0.894 | 0.822/0.944 | 0.882/0.934 | 0.844/0.801 | 0.810/0.807 | 0.449/0.459 |
| 8B6V_A/8B6W_A | 0.655 | 0.964/0.669 | 0.925/0.626 | 0.973/0.707 | 0.956/0.750 | 0.950/0.758 | 0.969/0.706 | 0.362/0.359 |
| 8DKF_L/8DOW_D | 0.764 | 0.943/0.955 | 0.982/0.952 | 0.981/0.962 | 0.972/0.957 | 0.979/0.943 | 0.984/0.957 | 0.916/0.889 |
| 7WKP_A/7WWU_I | 0.723 | 0.921/0.695 | 0.925/0.701 | 0.950/0.730 | 0.926/0.724 | 0.929/0.705 | 0.937/0.699 | 0.925/0.689 |
| 7PIS_9/7PIT_9 | 0.872 | 0.721/0.716 | 0.889/0.910 | 0.868/0.916 | 0.885/0.909 | 0.878/0.921 | 0.871/0.911 | 0.290/0.297 |
| 8DKE_P/8DKI_P | 0.813 | 0.963/0.963 | 0.862/0.969 | 0.949/0.941 | 0.953/0.960 | 0.963/0.945 | 0.947/0.968 | 0.960/0.855 |
| 8G4C_D/8G4D_D | 0.808 | 0.856/0.841 | 0.887/0.859 | 0.916/0.883 | 0.879/0.847 | 0.845/0.787 | 0.864/0.829 | 0.672/0.630 |
| 7ZF5_D/7ZF6_H | 0.801 | 0.956/0.937 | 0.977/0.923 | 0.924/0.944 | 0.972/0.901 | 0.928/0.933 | 0.910/0.950 | 0.676/0.620 |
| 8HKX_AS7P/8HKY_AS7P | 0.755 | 0.797/0.771 | 0.797/0.775 | 0.803/0.796 | 0.800/0.788 | 0.782/0.748 | 0.790/0.763 | 0.782/0.755 |
| 8SQZ_C/8SRM_C | 0.754 | 0.687/0.862 | 0.676/0.856 | 0.684/0.846 | 0.709/0.859 | 0.691/0.862 | 0.689/0.864 | 0.682/0.855 |
| 8HKX_S13P/8HKY_S13P | 0.769 | 0.786/0.768 | 0.795/0.785 | 0.797/0.772 | 0.801/0.766 | 0.788/0.759 | 0.782/0.770 | 0.741/0.731 |
| 8V2D_a/8V3B_A | 0.874 | 0.826/0.860 | 0.851/0.899 | 0.817/0.813 | 0.944/0.907 | 0.835/0.863 | 0.829/0.861 | N/A |
| 8EE5_H/8EF3_A | 0.846 | 0.824/0.681 | 0.928/0.865 | 0.936/0.908 | 0.966/0.956 | 0.866/0.812 | 0.844/0.721 | 0.838/0.778 |
| 8HC3_E/8HC6_L | 0.853 | 0.898/0.889 | 0.914/0.905 | 0.916/0.917 | 0.934/0.932 | 0.898/0.905 | 0.856/0.827 | 0.893/0.865 |
| 7QD7_AAA/7QH9_AAA | 0.900 | 0.985/0.957 | 0.982/0.964 | 0.985/0.981 | 0.984/0.973 | 0.977/0.960 | 0.983/0.954 | 0.305/0.306 |
| 7S8G_L/7UDS_L | 0.856 | 0.866/0.923 | 0.879/0.937 | 0.935/0.947 | 0.975/0.962 | 0.736/0.676 | 0.773/0.700 | 0.827/0.895 |
| 8I6O_B/8I6Q_B | 0.862 | 0.895/0.916 | 0.906/0.920 | 0.943/0.925 | 0.928/0.920 | 0.908/0.918 | 0.908/0.915 | 0.915/0.916 |
| 8HC3_H/8HC5_H | 0.878 | 0.868/0.840 | 0.892/0.933 | 0.909/0.912 | 0.875/0.894 | 0.819/0.775 | 0.856/0.865 | 0.634/0.668 |
| Mean-of-Best | | 0.796/0.784 | 0.825/0.817 | 0.832/0.829 | 0.837/0.834 | 0.801/0.785 | 0.803/0.793 | 0.654/0.659 |
| Median-of-Best | | 0.861/0.824 | 0.888/0.862 | 0.912/0.907 | 0.899/0.885 | 0.845/0.794 | 0.856/0.817 | 0.677/0.707 |

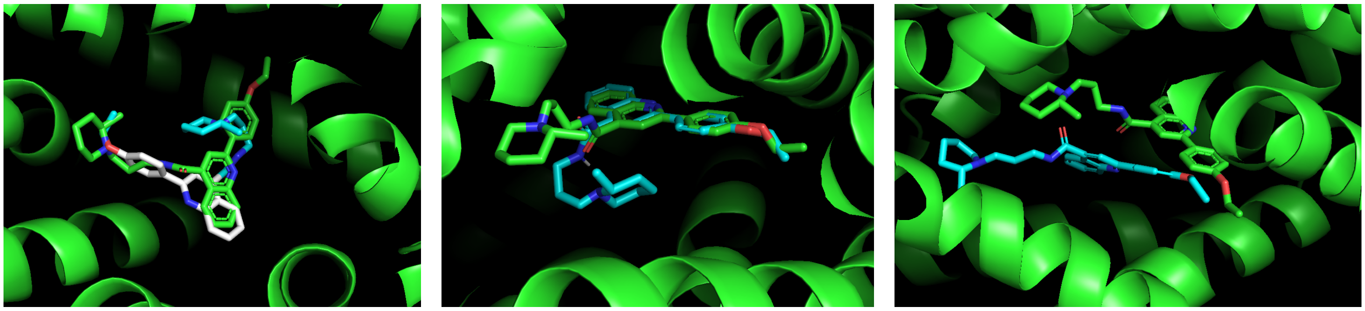

(a) QuickVina-W (b) Smina (c) Diffdock

Figure S2: Binding sites from three docking methods: (a) QuickVina-W (b) Smina (c) Diffdock. Green: Experimental strucuture of 8JZX and its binding ligand; blue: Docked ligand structure for different docking methods.

#### B.4 Structure Module on Diffused Inputs

In our early experiments, we tested the performance of AlphaFold with changed inputs to the structure module before fine-tuning. Surprisingly, replacing the zero-initialized structures with diffused structures did not affect the model's overall performance. We posit that the weight-sharing mechanism contributes to this robustness, as it was previously used to refine all zero-initialized and half-generated structures.

Specifically, we diffused the conformation of 4AKE with different diffusion times $t = (0.0, 0.1, \cdots, 1.0)$, and inputted these diffused structures into the structure module to generate predictions. From the metrics and visualized results reported in Fig. S3, the structure module generates conformations from diffused inputs with good qualities. Interestingly, the generated conformations are in-between the apo and holo states of the protein. Even when the un-diffused structure was inputted ($t = 0$), the model bent it slightly towards the conformation of 1AKE.

#### B.5 Details on Langevin Dynamics

We hereby specify the implementation of the overdamped Langevin dynamics. Specifically, we implement each step of the overdamped Langevin by taking a step in the reverse-time SDE (Eq. (4)) and then a step (with the same size) in the forward SDE (Eq. (3)). Specifically, the reverse-time SDE was discretized following Eq. (S11) - (S12), and the discretization of the forward SDE was introduced in a corresponding form as

$$\boldsymbol{c}^{(t)} = \sqrt{1 - \beta_{\text{pos}}^{(t,t-\tau)}} \boldsymbol{c}^{(t-\tau)} + \beta_{\text{pos}}^{(t,t-\tau)} \boldsymbol{\epsilon}_{\text{pos}}, \quad \boldsymbol{\epsilon}_{\text{pos}} \sim \mathcal{N}(\boldsymbol{0}, \boldsymbol{I}_3) \tag{S15}$$

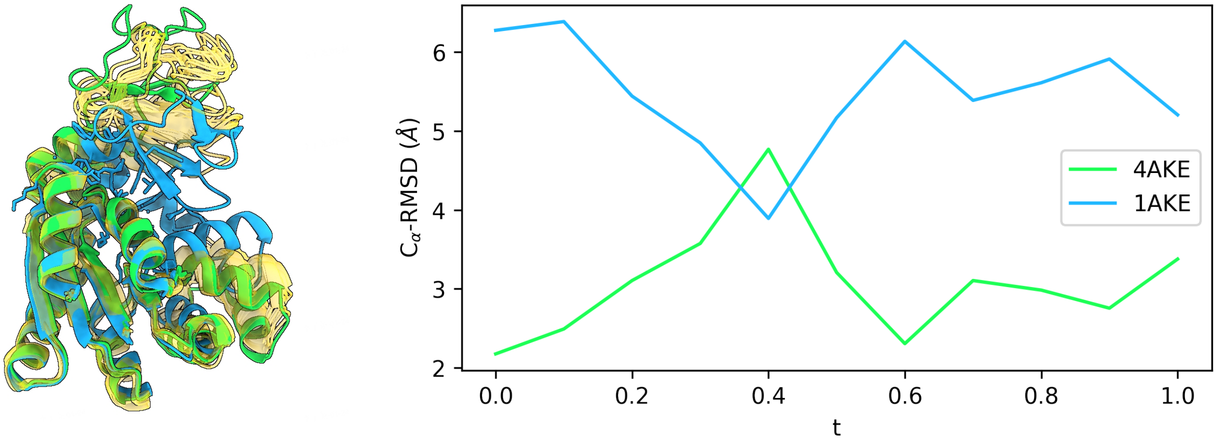


(a) Generated conformations. (b) RMSD to 4AKE and 1AKE.

Figure S3: AlphaFold behaves surprisingly well on diffused structures even without fine-tuning. In (a), green: 4AKE; blue: 1AKE; yellow: generated conformations from AlphaFold's structure module with diffused 4AKE with $t = (0.0, 0.1, \cdots, 1.0)$.

$$\boldsymbol{R}^{(t)} = \text{Exp}\left(\boldsymbol{\epsilon}_{\text{rot}}^{(t-\tau)}\right) \cdot \boldsymbol{R}^{(t-\tau)}, \quad \boldsymbol{\epsilon}_{\text{rot}}^{(t-\tau)} \sim \text{IGSO3}\left(\beta_{\text{rot}}^{(t,t-\tau)}\right). \tag{S16}$$

The implementation can be justified in two ways: 1) by observing that Eq.(19) is formally equivalent to the sum of Eq.(3) and Eq.(4); 2) by observing that this implementation does not change the marginal distributions of $\mathcal{P}$, therefore providing a sampling method of $p(\mathcal{P}, t)$.

Notably, when analyzing the generated trajectories, we are more focused on the transition and distribution of the per-step predictions $\hat{\mathcal{P}}_0^{(t)}$, which have correct geometries and *conform to a distribution induced by the model* $m_\theta$ *from* $p(\mathcal{P}^{(t)}, t)$. Under condition $t \to 0$, as is required in,[10] the two spaces tend to be equivalent. However, drastic changes of conformation can hardly be observed when $t \to 0$, because the diffused structure $\mathcal{P}^{(t)}$ contains too much information and the model would be trapped into the local minimum around $\mathcal{P}^{(t)}$.

Besides Fig. 5(e-f), we also plot the 2D projection of trajectories in terms of RMSD to two conformations as Fig. S4. Notably, the trajectory well covers a bridge between 1AKE and 4AKE with continuity.

#### B.6 More Visualization Results

We visualize the inference process on 1AKE case by UFConf, as shown in Fig. S5. A balanced sampling between 1AKE and 4AKE is observed. The diffused conformations, $\mathcal{P}^{(t)}$, converge gradually to

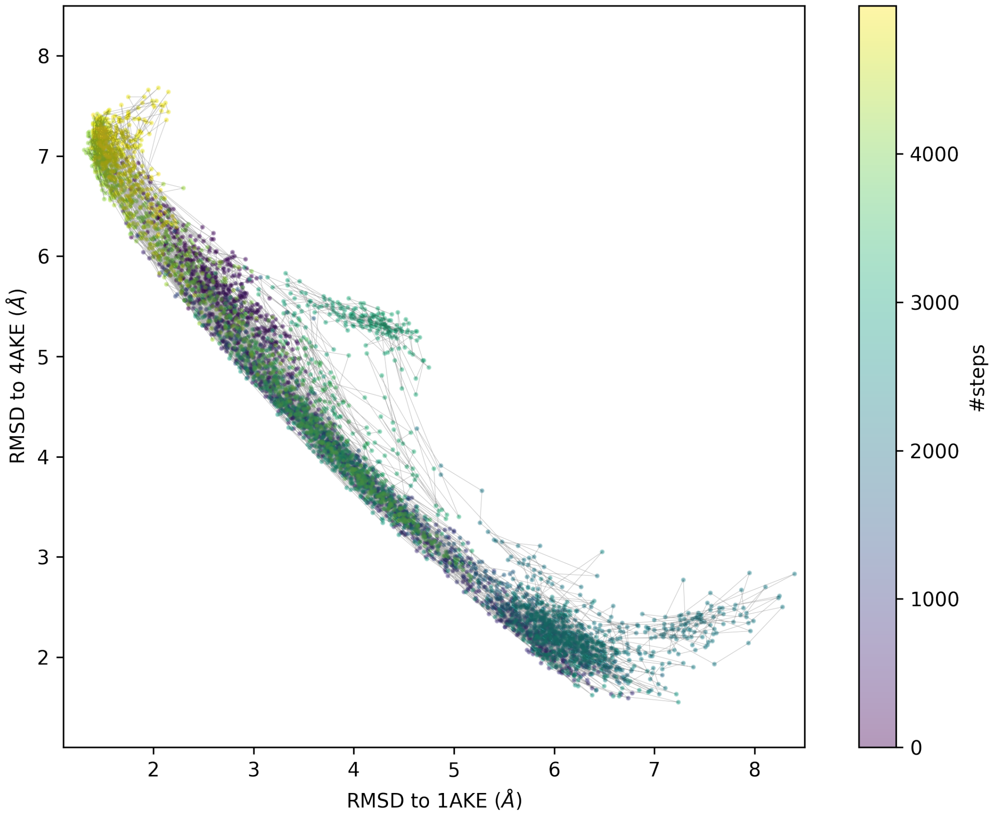


Figure S4: Trajectory of 5000 steps of Langevin dynamics

.

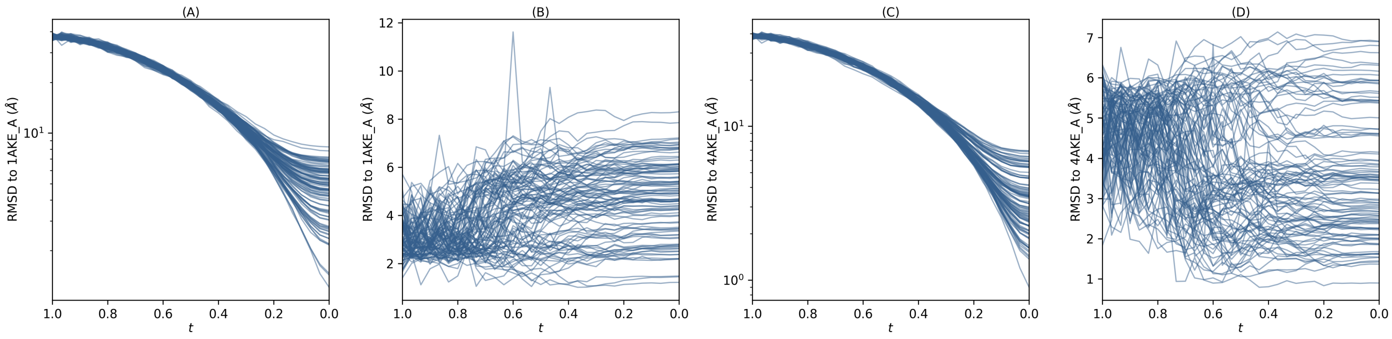


Figure S5: Per-step RMSDs to 1AKE / 4AKE in the inference process.(A) RMSDs of noisy structures versus 1AKE; (B) RMSDs of predicted structures versus 1AKE; (C) RMSDs of noisy structures versus 4AKE; (D) RMSDs of predicted structures versus 4AKE.

conformations with correct geometries. The predicted conformations, $\hat{\mathcal{P}}^{(t)}$, retain correct geometries all the time and are all close to real conformations. Global folding is determined during the early stage of inference, while the local precision is enhanced (not embodied in the figure) during the late half.

We also visualize the interpolated structures from 1AKE to 4AKE in Fig. S6. To make the figure clear we only select 5 intermediate structures along the pathway, from the closest to the 1AKE to the closest to the 4AKE.

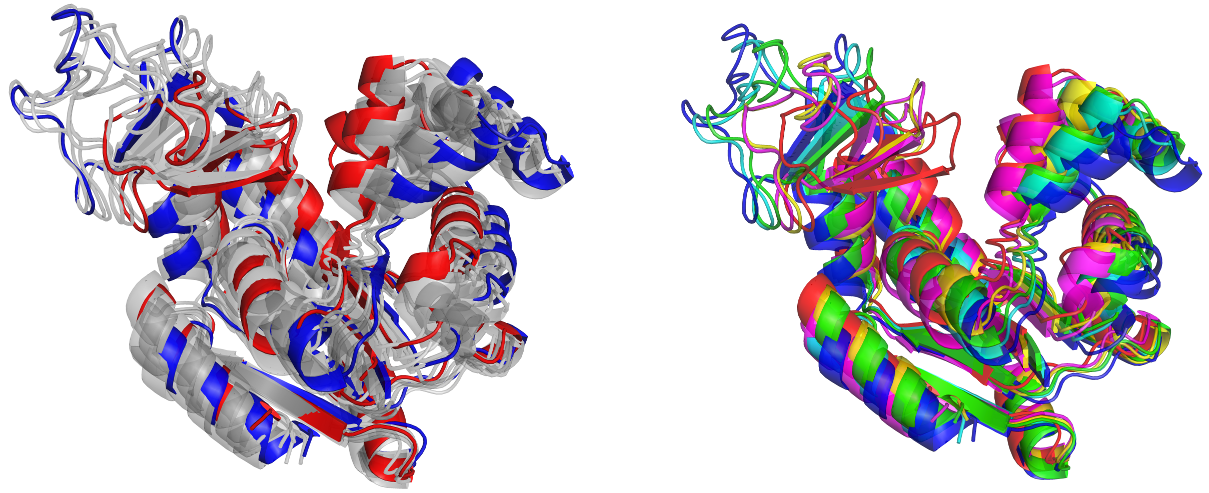

(a) (b)

Figure S6: Interpolated structures from 1AKE to 4AKE. (a) Red: 1AKE structure. Blue: 4AKE structure. Gray: generated interpolated structures by UFConf. (b)Generated interpolated structures by UFConf, the structures are colored red, magnenta, yellow, cyan and blue from 1AKE to 4AKE.

To see the stability of different generated conformations, we visualize the generated conformations for the 8D01_L/8D0Y_L system from MSA-subsampling and the 8I6O_B/8I6Q_B system from AF-cluster methods, as shown in Fig. S7. We can see from the figure that the generated conformations with low TM-score (Fig. S7(b) and (d)) against the experimental structures have fewer secondary structures compared with the conformations with high TM-score (Fig. S7(a) and (c))against the experimental structures. This is clear for the 8I6O_B/8I6Q_B system as the conformation in Fig. S7(d) has much fewer beta sheet than Fig. S7(c). For the 8D01_L/8D0Y_L system, the conformation in Fig. S7(b) has incomplete beta sheet in the lower right of the figure compared with Fig. S7(a).

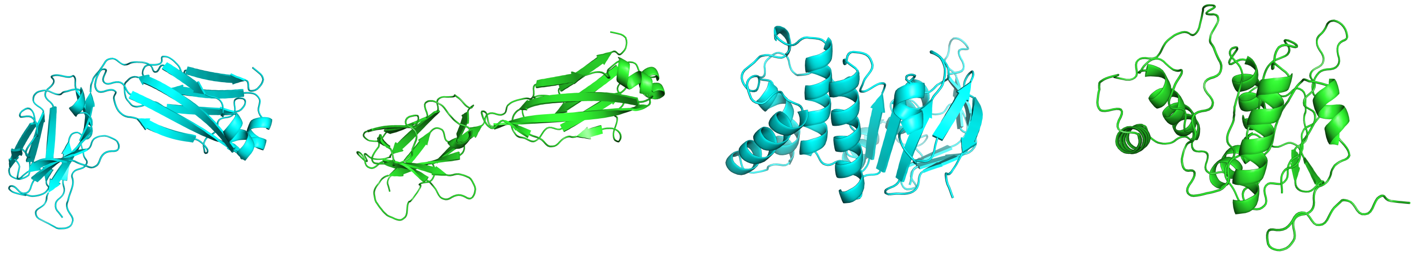

(a) (b) (c) (d)

Figure S7: (a) Generated conformation with high TM-score against 8D01_L/8D0Y_L structures by MSA-subsampling; (b) Generated conformation with low TM-score against 8D01_L/8D0Y_L structures by MSA-subsampling; (c) Generated conformation with high TM-score against 8I6O_B/8I6Q_B structures by AF-cluster; (d) Generated conformation with low TM-score against 8I6O_B/8I6Q_B structures by AF-cluster.

For all conformations generated for RAC-47 from UFConf, we visualize them in Fig. S8-Fig. S11.

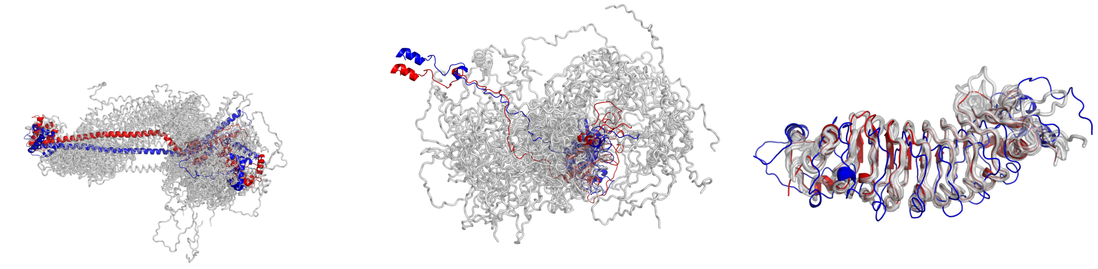

(a) 7Z3N_C/7Z3O_C (b) 7R5J_F0/7R5K_F0 (c) 7YCO_B/8H6F_X

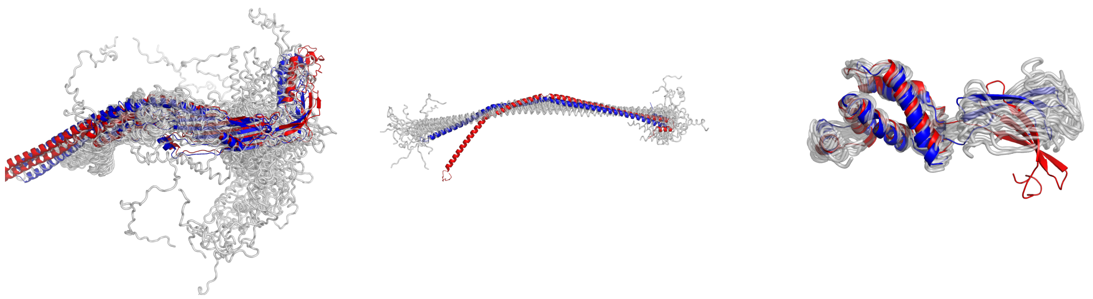

(d) 8TOC_b/8TV9_a (e) 8TH8_S/8TID_s (f) 7P37_A/7P3F_A

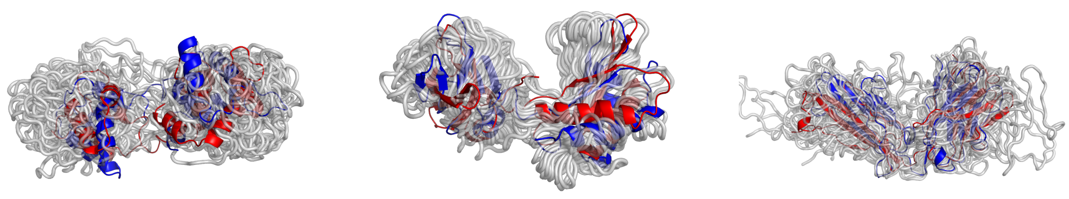

(g) 7UTI_Z/7UTL_d (h) 8BDV_A/8BH7_A (i) 7SJO_H/7SJP_H

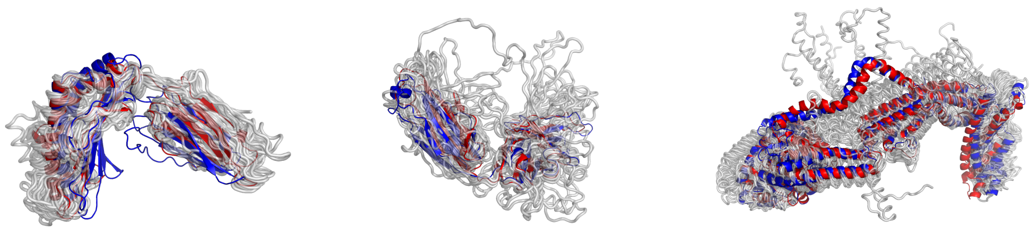

(j) 7TUC_A/7TUE_A (k) 8DKF_H/8DOW_C (l) 8EFR_A/8EFT_A

Figure S8: Comparative visualization of PDB in **RAC-47** benchmark (Part 1)

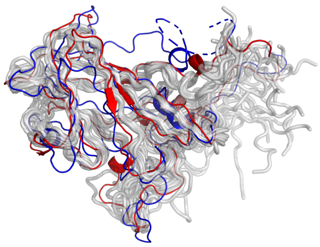

(a) 8E2Y_A/8E31_A

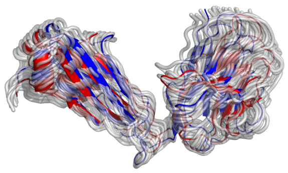

(b) 8D01_L/8D0Y_L

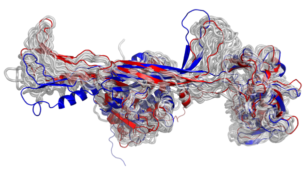

(c) 8BFL_A/8BFP_A

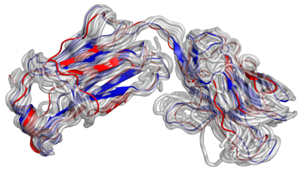

(d) 7SJO_F/7SJP_L

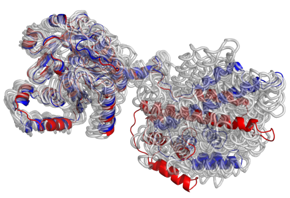

(e) 8DUE_A/8DVF_A

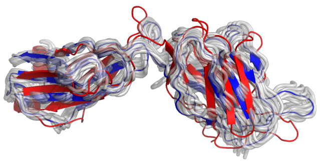

(f) 7ZWM_E/7ZXF_E

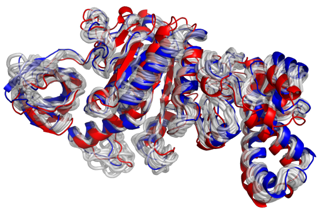

(g) 7Y5A_D/7Y5B_D

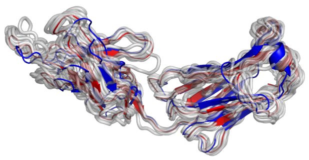

(h) 8FWF_L/8FYM_L

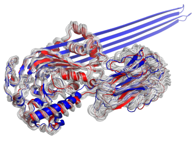

(i) 8B6V_A/8B6W_A

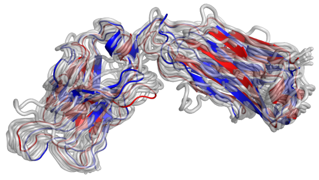

(j) 8DKF_L/8DOW_D

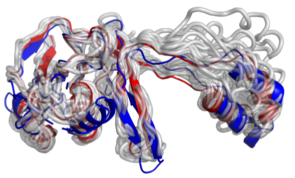

(k) 7WKP_A/7WWU_I

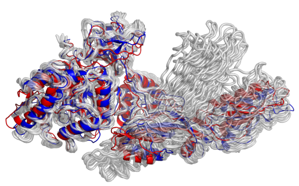

(l) 7PIS_9/7PIT_9

Figure S9: Comparative visualization of PDB in **RAC-47** benchmark (Part 2)

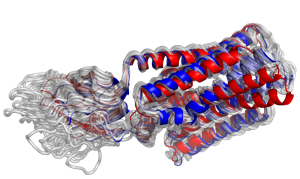

(a) 8DKE_P/8DKI_P

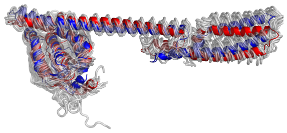

(b) 8G4C_D/8G4D_D

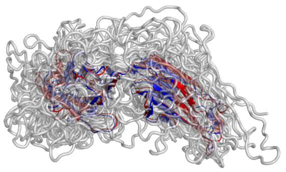

(c) 7ZF5_D/7ZF6_H

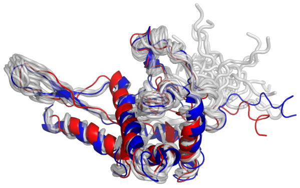

(d) 8HKX_AS7P/8HKY_AS7P

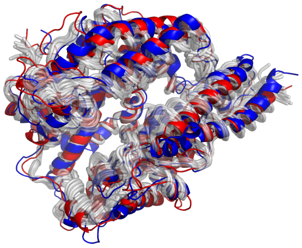

(e) 8HFX_E/8IFY_E

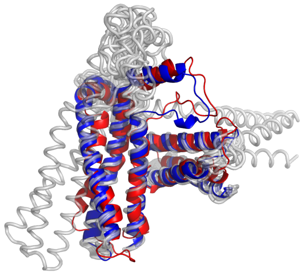

(f) 8SQZ_C/8SRM_C

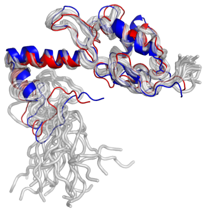

(g) 8HKX_S13P/8HKY_S13P

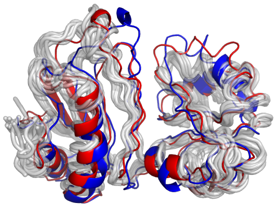

(h) 8HKY_AL1P/8HKZ_AL1P

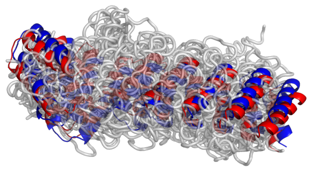

(i) 8V2D_a/8V3B_A

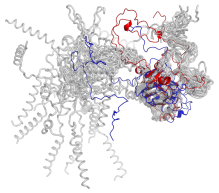

(j) 8E39_A/8E3A_A

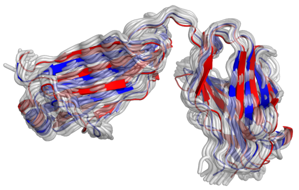

(k) 8TCA_L/8VEV_B

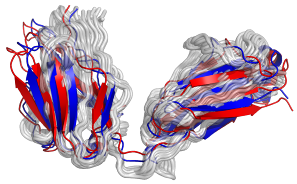

(l) 8EE5_L/8EF3_B

Figure S10: Comparative visualization of PDB in **RAC-47** benchmark (Part 3)

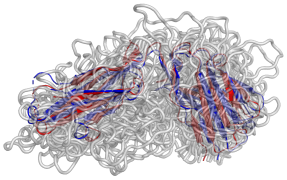

(a) 8EE5_H/8EF3_A

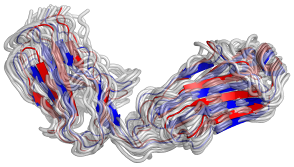

(b) 7ZF5_F/7ZF6_L

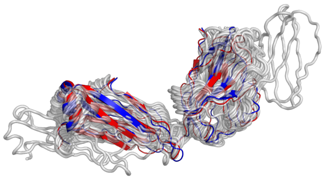

(c) 8HC3_E/8HC6_L

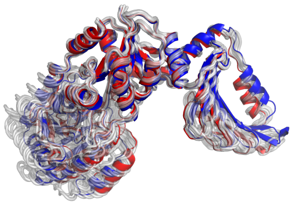

(d) 7QD7_AAA/7QH9_AAA

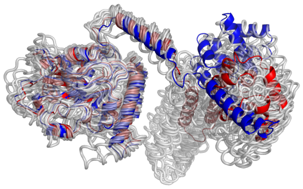

(e) 7T22_A/7T23_B

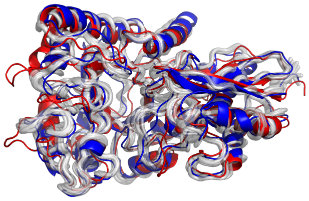

(f) 8UOP_A/8UOY_B

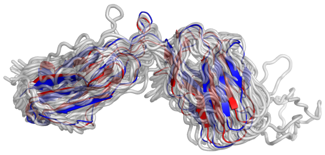

(g) 7S8G_L/7UDS_L

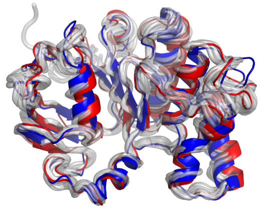

(h) 8I6O_B/8I6Q_B

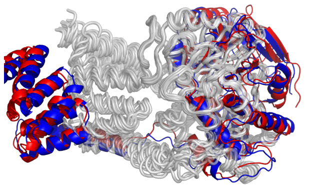

(i) 8GAE_E/8GFT_E

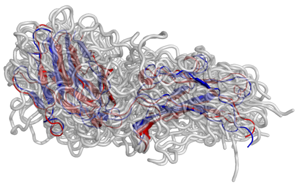

(j) 8HC3_H/8HC5_H

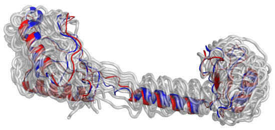

(k) 8JSG_A/8JSH_A

Figure S11: Comparative visualization of PDB in **RAC-47** benchmark (Part 4)

### References


(1) Steinegger, M.; Söding, J. MMseqs2 enables sensitive protein sequence searching for the analysis of massive data sets. *Nature biotechnology* **2017**, *35*, 1026–1028.

(2) Zhang, Y.; Skolnick, J. TM-align: a protein structure alignment algorithm based on the TM-score. *Nucleic acids research* **2005**, *33*, 2302–2309.

(3) van Kempen, M.; Kim, S. S.; Tumescheit, C.; Mirdita, M.; Gilchrist, C. L.; Söding, J.; Steinegger, M. Foldseek: fast and accurate protein structure search. *Biorxiv* **2022**, 2022–02.

(4) Blondel, V. D.; Guillaume, J.-L.; Lambiotte, R.; Lefebvre, E. Fast unfolding of communities in large networks. *Journal of statistical mechanics: theory and experiment* **2008**, *2008*, P10008.

(5) Ho, J.; Jain, A.; Abbeel, P. Denoising diffusion probabilistic models. *Advances in neural information processing systems* **2020**, *33*, 6840–6851.

(6) Song, Y.; Sohl-Dickstein, J.; Kingma, D. P.; Kumar, A.; Ermon, S.; Poole, B. Score-based generative modeling through stochastic differential equations. *arXiv preprint arXiv:2011.13456* **2020**,

(7) Li, Z.; Liu, X.; Chen, W.; Shen, F.; Bi, H.; Ke, G.; Zhang, L. Uni-Fold: an open-source platform for developing protein folding models beyond AlphaFold. *bioRxiv* **2022**, 2022–08.

(8) Berman, H. M.; Westbrook, J.; Feng, Z.; Gilliland, G.; Bhat, T. N.; Weissig, H.; Shindyalov, I. N.; Bourne, P. E. The protein data bank. *Nucleic acids research* **2000**, *28*, 235–242.

(9) DeLano, W. L.; others Pymol: An open-source molecular graphics tool. *CCP4 Newsl. Protein Crystallogr* **2002**, *40*, 82–92.

(10) Arts, M.; Garcia Satorras, V.; Huang, C.-W.; Zügner, D.; Federici, M.; Clementi, C.; Noé, F.; Pinsler, R.; van den Berg, R. Two for one: Diffusion models and force fields for coarse-grained molecular dynamics. *Journal of Chemical Theory and Computation* **2023**, *19*, 6151–6159.
